# Functional characterization of RELN missense mutations involved in recessive and dominant forms of Neuronal Migration Disorders

**DOI:** 10.1101/2021.05.25.445586

**Authors:** Martina Riva, Sofia Ferreira, Vera P. Medvedeva, Frédéric Causeret, Olivia J. Henry, Charles-Joris Roux, Céline Bellesme, Elena Freri, Elena Parrini, Dragana Josifova, Renzo Guerrini, Nadia Bahi-Buisson, Alessandra Pierani

**Author notes:** equal contribution (Renzo Guerrini, Nadia Bahi-Buisson).

## Abstract

RELN is a large secreted glycoprotein that acts at multiple steps of cerebral cortex development, including neuronal migration. Only recessive mutations of the Reelin gene (*RELN*) have been associated with human cortical malformations and none has been functionally characterized. We identified novel missense *RELN* mutations in both compound and *de novo* heterozygous patients exhibiting an array of neuronal migration disorders (NMDs) as diverse as pachygyria, polymicrogyria and heterotopia. Most mutations caused defective RELN secretion *in vitro* and, when ectopically expressed in the embryonic mouse cortex, affected neuronal aggregation and/or migration *in vivo*. We determined the *de novo* heterozygous mutations acted as dominant negative and demonstrated that *RELN* mutations mediate not only recessive, but also dominant NMDs. This work assesses for the first time the pathogenicity of RELN mutations showing a strong genotype-phenotype correlation. In particular, the behavior of the mutant proteins *in vitro* and *in vivo* predicts the severity of cortical malformations and provides valuable insight into the pathogenesis of these disorders.

## INTRODUCTION

Abnormal brain development participates in the pathophysiology of multiple neurodevelopmental disorders. The neocortex is composed of six layers that are built during embryonic development through highly orchestrated processes of successive generation of cohorts of glutamatergic neurons in the proliferative zones and their radial migration to form distinct layers (*1*). The inside-out sequence in the formation of these layers, whereby later-born neurons bypass earlier-born ones to position more superficially, is a unique characteristic of the mammalian neocortex and was shown to be driven by the Reelin (RELN) protein (*2, 3*). RELN is an extracellular matrix (ECM) glycoprotein, which is cleaved in the extracellular environment at two main specific sites, between repeats 2-3 (N-t site) and repeats 6-7 (C-t site) (*4-7*), by cleaving enzymes such as matrix metalloproteinases (*5, 8-10*). Studies on RELN proteolysis have identified an N-terminal (N-t) region necessary for the extracellular homodimerization of the protein (*11, 12*), a central region containing the binding sites (R3-6) for the two best known RELN receptors (apolipoprotein E receptor 2 (ApoER2) and very low density lipoprotein receptor (VLDLR)), (*4, 13-15*) and a C-terminal (C-t) region required for efficient activation of downstream signaling (*16*), but whose importance for secretion is still debated (*17, 18*). The full-length protein is generally more efficient in activating the transduction cascade probably due to the N-terminal region that promotes homodimerization through disulfide linkage, and the central region that mediates proper folding (*11, 12, 19-21*). Although RELN has been studied for the last 30 years, its functions are still unclear. On one hand, it is proposed to act as an attractant cue (*22*), and on the other hand it is thought to serve as a “detach and go” signal instructing the migrating neurons to desengage from the radial glia and switch from locomotion mode of migration to terminal somal translocation stopping below the marginal zone (*4, 23-28*). RELN has been initially studied via the characterization of the *reeler* (*rl/rl*) homozygous mouse mutant (*3, 29*), which shows, amongst a large panel of phenotypes, a profound disorganization of cortical lamination, largely due to an impairment in migration of pyramidal neurons (*2, 30*). On the contrary, heterozygous *reeler* (*rl/+*) mice (haploinsufficient for RELN) show no defects in cortical layering, although exhibit a spectrum of cognitive and behavioral abnormalities (*31, 32*), emphasizing the relevance of RELN expression levels in higher brain functions.

In humans, recessive (homozygous or compound) *RELN* mutations have been associated to different patterns of lissencephaly with cerebellar hypoplasia (LCH), a profound developmentally disabling disease (*33-35*), and suffering of epilepsy. A total of ten pathogenic *RELN* variants in eight families were identified. Five families carried homozygous loss-of-function mutations due to balanced reciprocal translocation (*34*), disruption of splicing (*33*), nonsense or frameshift (*35*). Three families bore compound heterozygous missense and/or truncating mutations (*35, 36*). In addition, one single patient with a history of polymicrogyria, microcephaly and epilepsy was described with two missense biallelic mutations (*37*). Recently, several heterozygous *RELN* mutations were identified as risk factors for multiple neuropsychiatric and neurodegenerative disorders, such as schizophrenia, bipolar disorders, Autism Spectrum Disorders (ASD) and Alzheimer’s disease (*38-40*). In addition, heterozygous *RELN* mutations account for 17.5% of familial cases of autosomal dominant lateral temporal lobe epilepsy (ADLTE) with relatively low penetrance (*41*). They are mainly missense variants, which result in changes of structurally important amino acids suggested to perturb protein folding (*41*). However, only one *RELN* missense mutation associated with ASD thus far has been functionally characterized *in vitro* (*42*) and the mechanisms underlying ASD and ADTLE remain undetermined. Furthermore, it is unknown whether the phenotypes arise from hemizygous gain-of-function (GOF) or loss-of-function (LOF) and, importantly, which specific sub-function of RELN may be affected to cause such high variety of pathologies.

Here we report 6 patients with compound (C), maternally-inherited (MI) and *de novo* (DN) heterozygous *RELN* missense mutations associated with a spectrum of malformations of cortical development (MCDs), namely pachygyria (reduced cortical gyration with shallow sulci and broad gyri) or polymicrogyria (excessive number of abnormally small cortical gyri) (*43*) and, for the first time, in the absence of cerebellar hypoplasia. We functionally characterized each mutation through a set of *in vitro* and *in vivo* assays to assess the secretion and processing of the mutated proteins and their capacity to form aggregates and regulate neuronal migration upon their ectopic expression in the mouse cerebral cortex. We assessed their pathogenicity demonstrating that all mutations alter at least one of the studied processes and the extent of that interference correlate with the severity of the pathology. Moreover, we provide the first evidence that heterozygous *de novo RELN* mutations can cause autosomal dominant NMDs. Our findings indicate that the pathological defect of RELN secretion and function contributes to NMDs risk, shedding light on the involvement of RELN in the etiology of MCDs.

## RESULTS

### Cortical malformations in patients carrying *RELN* variants

We identified six children with cortical malformations without LCH harboring *RELN* (NM_005045.4) missense variants (Fig. 1). Two were compound heterozygous (C1, C2) and four were heterozygous, with two *de novo* (DN1, DN2), and two brothers bearing the same maternally-inherited mutation (MI1/2). Affected children were diagnosed at 1-8 years of age with hypotonia and cognitive developmental delays. The first patient (C1) displayed bilateral fronto-temporo-parietal polymicrogyria and periventricular nodular heterotopia at brain magnetic resonance imaging (MRI) (Fig. 1a). A dedicated Next Generation Sequencing (NGS) panel including genes associated with MCDs revealed the c.5461T>C/c.3839G>A (p.Tyr1821His/p.Gly1280Glu) compound heterozygous *RELN* mutations (Fig. 1b,c). The second patient (C2) showed bilateral pachygyria most prominent on frontal regions (Fig. 1a) and harbored the c.1949T>G/c.1667A>T (p.Ile650Ser/p.Asp556Val) compound heterozygous *RELN* mutations at the NGS analysis of a panel for MCDs genes (Fig. 1b,c). Patients MI1 and MI2, two brothers (hence referred as MI1/2), presented MRI imaging consistent with bilateral perisylvian polymicrogyria (Fig. 1a). In these two patients, an NGS panel for genes associated with MCDs and intellectual disability revealed the c.2737C>T (p.Arg913Cys) missense substitution in the *RELN* gene (Fig. 1b,c), which they both inherited from their apparently healthy, but unexamined, mother. In the context of clinical findings and available evidence, the substitution identified in this family was classified as a variant of uncertain significance (VUS). No further variants of significance were identified by whole-exome sequencing (WES) in these brothers. The last two patients, hereafter DN1 and DN2, presented at the MRI pachygyria with severe or moderate simplified gyral pattern (Fig. 1a), respectively. NGS analysis of a panel for MCDs genes revealed a c.1615T>C (p.Cys539Arg) *de novo* mutation in patient DN1 and a c.9619C>T (p.Arg3207Cys) in DN2 (Fig. 1b,c). All patients had normal comparative genomic hybridization array (CGH-Array). At the exception of MI1/2, all patients were born from non-consanguineous healthy parents. Among all patients, only M1 suffered of epilepsy.

**Fig. 1.**
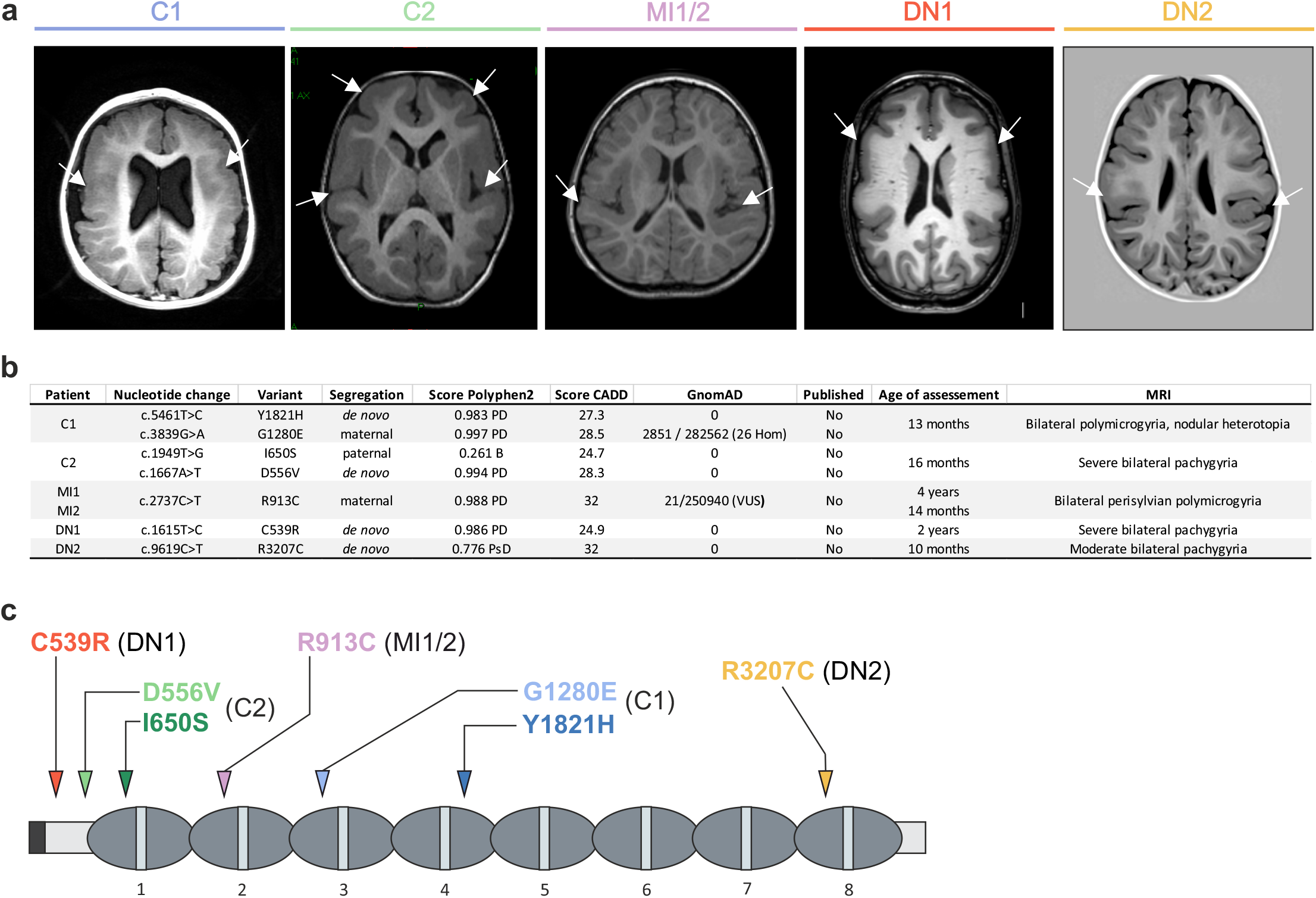
Cortical malformations in heterozygous patients associated with *RELN* missense variants. **a** MRIs from compound (C), maternal inherited (MI) and *de novo* (DN) patients. C1 exhibits bilateral fronto-parietal polymicrogyria with nodular heterotopia, C2 frontal predominant bilateral pachygyria, MI1/2 bilateral perisylvian polymicrogyria, DN1 and DN2 frontal predominant bilateral pachygyria, severe and moderate respectively. Representative axial T1 section of the cortical malformation (white arrows). **b** Recapitulative table of patients’ phenotype and genotype with inheritance and pathogenicity score. PD: probably damaging, B: benign, PsD: possibly damaging, CADD: combined annotation dependent depletion, GnomAD: genome aggregation database, Hom: homozygous, VUS: variant of uncertain significance. **c** Primary structure of the RELN protein. Arrows indicate the position of missense mutations, each color corresponds to one patient (C1 in blue and C2 in green, MI1/2 in pink, DN1 in red and DN2 in yellow).

These results suggest that heterozygous *RELN* variants are associated with a variety of cortical malformations, as diverse as those thought to raise from abnormal neuronal migration, such as pachygyria, or postmigrational development, such as polymicrogyria, and this in the absence of cerebellar hypoplasia, the hallmark of RELN-dependent autosomal recessive lissencephaly.

### *RELN* missense mutations affect its secretion and cleavage

Previously reported patients with LCH showed a decrease in serum levels of RELN (*33*) suggesting that the mutations resulted in a null allele. Moreover, the only up to now functional *in vitro* analysis of a missense *RELN* mutation associated with ASD described a reduction of protein secretion (*42*). We thus investigated whether the missense mutations identified in the 6 patients with MCDs could affect the expression and/or secretion of RELN. We introduced each of the seven point-mutations into the mouse *RELN* sequence (all of the affected residues being conserved but shifted +1aa compared to human, see Material and Methods). Plasmids carrying the mouse WT-RELN or the different mutations were transfected into HEK293T cells and RELN levels in the culture media and cell lysates were compared by immunoblotting using G10 antibodies recognizing epitopes in the N-terminal region (*44*) (Fig. 2).In mock- and GFP-transfected cells no signal was detected either in the cell lysate or culture media (data not shown). Upon WT-RELN transfection, a single full-length (FL) 450kDa band was observed in the cell fraction, whereas the FL 450 kDa, the two complementary fragments resulting from the N-t cleavage NR2 (150 kDa) and R3-8 (250 kDa), and those resulting from the C-t cleavage NR6 (340 kDa) and R7-8 (80 kDa), were visible in the secreted fraction (Fig. 2a,c and Supplementary Fig. 1), indicating that WT-RELN is efficiently secreted and processed as expected (*45*). In the cell fraction, significantly increased levels of FL RELN (450 kDa) were observed for D556V, C539R and R3207C transfected cells compared to WT (Fig. 2b)., In the secreted fraction, the total amount of Y1821H mutant was increased. In contrast, we observed a 47% and 65% decrease of RELN in the media of I650S- and of D556V-transfected cells, respectively. An even stronger effect was observed for the C539R and R3207C mutations for which both FL and all RELN proteolytic fragments were barely detectable in the culture media (Fig. 2c), even when the lanes were deliberately overexposed (not shown). Similar changes in total secreted RELN caused by the different mutations were detected by immunoblotting with the 12/14 antibodies that recognize the C-terminal region of the protein (Supplementary Fig. 1a,b). To investigate how RELN cleavage was occurring we also analyzed the relative amount of each fragment compared to the total amount of secreted protein for each mutation using the G10 antibodies. Amongst the mutations that produced detectable extracellular RELN, exclusively the D556V mutation displayed a 4-fold reduction of NR6 (Fig. 2d and Supplementary Fig.1c), thus indicating an impairment in the C-t cleavage process.

**Fig. 2.**
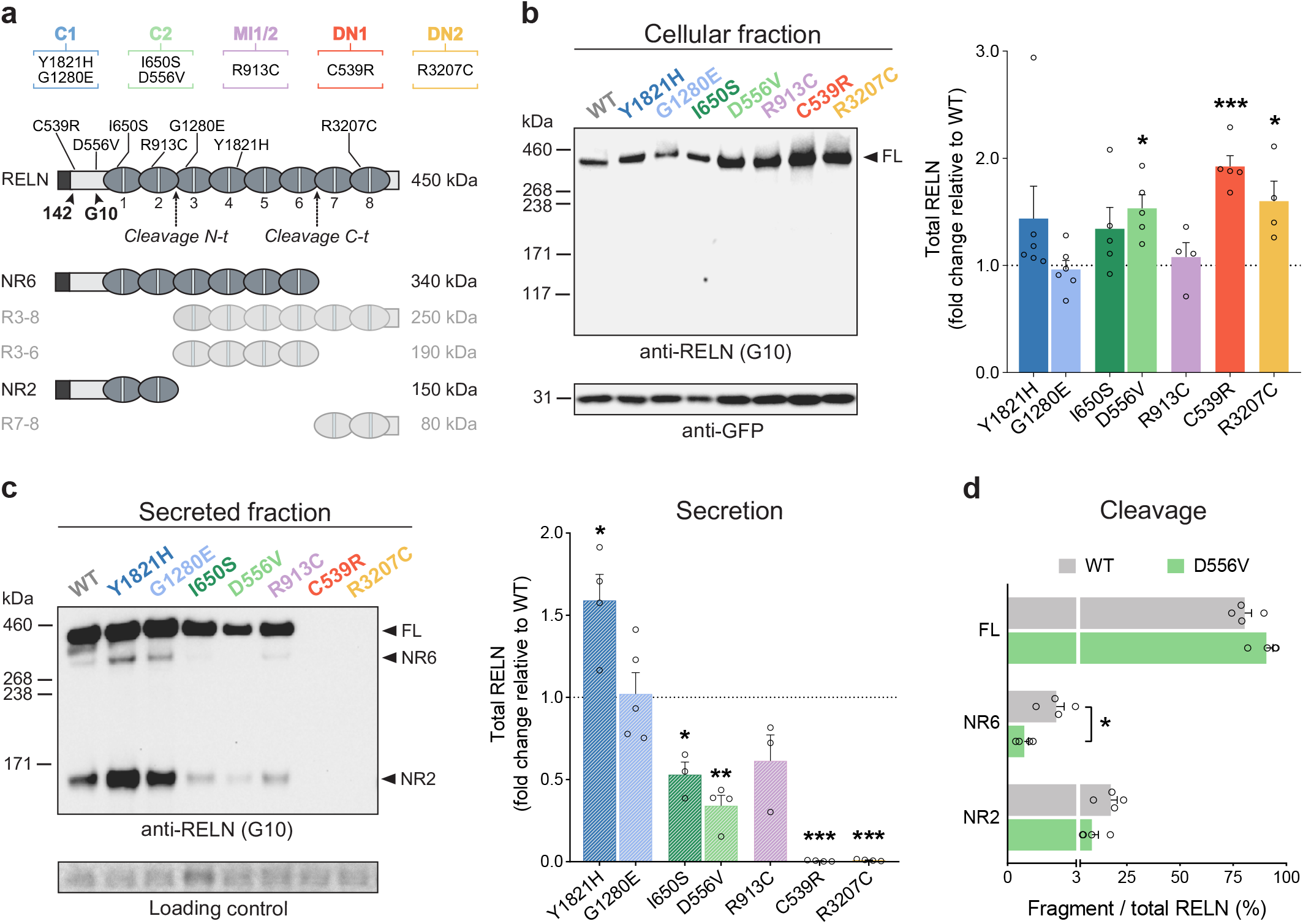
Missense mutations alter secretion and proteolytic processing of RELN. **a** Schematic of the full-length RELN protein (450 kDa), its cleavage sites N-t and C-t (dotted arrows), and its five cleaved products (NR6, R3-8, R3-6, NR2, R7-8). The binding region of the 142 and G10 antibodies and the position of *RELN* mutations are indicated with black arrowheads and lines, respectively. Patient colour coding and corresponding mutations are indicated above. **b** Representative immunoblotting of the cellular fraction of HEK293T cells transfected with either WT-RELN or mutants-RELN, probed with anti-RELN G10 or anti-GFP antibodies. Right panel shows the densitometric analysis of RELN-FL normalized to GFP signal (n=4-6 independent experiments). More RELN was detected in lysates of cells transfected with D556V (\**p*=0.0138), C539R (\*\*\**p*=0.0008) and R3207C (\**p*=0.0486) mutants compared to WT. **c** Representative immunoblotting of the secreted fraction of HEK293T cells transfected with either WT-RELN or mutants-RELN, probed with anti-RELN G10 antibody. Right panel shows the densitometric analysis of total RELN levels normalized to total protein (Ponceau S) (n=3-5 independent experiments). Higher levels of Y1821H mutant were detected in the media (\**p*=0.0334) whereas I650S-, D556V-, C539R- and R3207C-transfected cells secreted lower RELN levels (\**p*=0.0259, \*\**p*=0.0019, \*\*\**p*<0.0001 and \*\*\**p*<0.0001, respectively). Protein standard sizes (kDa) are indicated on the left side of the blots. **d** Densitometric analysis of the proportion (%) of RELN-FL, NR6 (from C-t cleavage) and NR2 (from N-t cleavage) relative to the total amount of secreted D556V protein or secreted WT (n=4; FL: *p*=0.0571; NR6: \**p*=0.0286; NR2: *p*=0.0571).

Taken together, these observations indicate that mutations in the *de novo* and compound C2 heterozygous patients cause severe and mild deficiency in RELN secretion, respectively, whereas the Y1821H mutation of patient C1 appears to enhance it. The significant accumulation of intracellular RELN detected for the D556V, C539R and R3207C mutations is consistent with their pronounced deficit in RELN secretion. Additionally, we show that specifically the D556V mutation of patient C2 affects cleavage of RELN in the extracellular media.

### *RELN de novo* heterozygous mutations behave as dominant negative forms *in vitro*

To assess how RELN generated from mutant alleles might influence the total RELN levels as in the genetic context of the patients, we modeled *in vitro* the compound heterozygous patients’ genotypes by co-transfection of Y1821H and G1280E mutations for C1, or I650S and D556V for C2, and those of the heterozygous patients by co-transfection of WT-RELN with either the R913C, C539R or R3207C mutations for patient MI1/2, DN1 and DN2, respectively.

Western blot analysis of C1 and MI1/2 mutations showed unchanged amounts of RELN in both the cellular and the secreted fraction (Fig. 3a,b), while C2 mutations displayed an increase in the intracellular levels (Fig. 3a) and a 47% reduction of total secreted RELN (Fig. 3b).

**Fig. 3.**
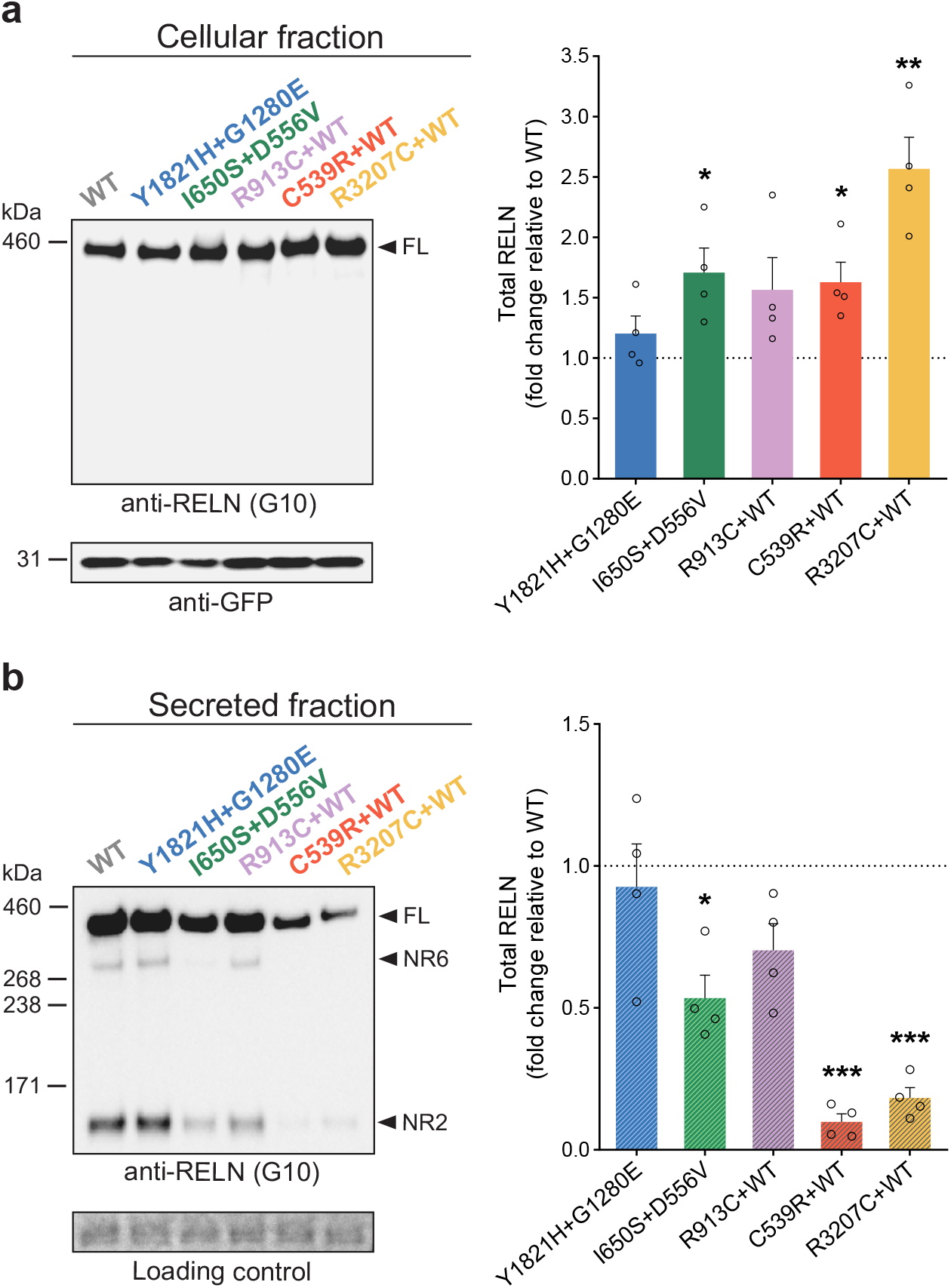
*De novo* heterozygous *RELN* mutations behave as dominant negative. **a** Representative immunoblotting of the cellular fraction of HEK293T cells co-transfected with Y1821H and G1280E mutations, or I650S and D556V mutations, or co-transfected with WT-RELN and R913C, C539R, or R3207C mutations, probed with anti-RELN G10 or anti-GFP antibodies. Right panel shows the densitometric analysis of RELN-FL normalized to GFP signal (n=4). Increased RELN levels were found in the conditions I650S+D556V (\**p*=0.0398), C539R+WT (\**p*=0.0325) and R3207C+WT (\*\**p*=0.0092). **b** Representative immunoblotting of the secreted fraction of co-transfected HEK293T cells, probed with anti-RELN G10 antibody. Right panel shows the densitometric analysis of total RELN levels normalized to total protein (Ponceau S) (n=4). Cells transfected with either I650S+D556V, C539R+WT and R3207C+WT presented decreased RELN levels in the media (\**p*=0.0104, \*\*\**p*<0.0001 and \*\*\**p*=0.0002) when compared to WT. Protein standard sizes (kDa) are indicated on the left side of the blots.

When WT-RELN was co-transfected with either C539R or R3207C mutant forms, RELN levels were strongly reduced in the secreted fraction (90% and 80%, respectively) (Fig. 3b) and in parallel the cell lysates displayed increased RELN protein compared to WT control (Fig. 3a). Similar differences were detected using the C-terminal antibodies 12/14 (Supplementary Fig. 2a). These results indicate that Y1821H and G1280E mutant proteins, when both present, are secreted as efficiently as WT proteins, whereas the co-existence of I650S and D556V mutations diminish RELN secretion. As for the heterozygous mutations, the MI1/2 mutant form is unlikely to interfere with the WT-RELN protein whereas the *de novo* C539R and R3207C strongly impair WT RELN secretion, demonstrating a dominant negative effect. To go further in the molecular mechanisms, we performed blots in non-reducing conditions to identify dimerized forms of RELN. As expected (*11*) in the secreted fraction a high proportion of the WT protein is present as a homodimer of around 900kDa (Supplementary Fig. 2b). As never described before, we were also able to observe these homodimers in the cellular fraction for all the conditions including the WT protein (Supplementary Fig. 2c), suggesting intracellular multimerization as a possible mechanism by which C539R and R3207C could be subtracting the WT protein from secretion.

### Reduced serum RELN levels of patient C2 correlate with in vitro observations

RELN is expressed in several tissues other than the brain, such as liver, pancreas and intestine (*46, 47*), and circulating RELN is detectable in the serum of adult mammals (*48*). Altered blood RELN levels was detected in several patients diagnosed with different cortical malformations when compared to control individuals (*33, 41, 49, 50*). RELN in the blood undergoes post-translational processing similarly to that observed in the brain. To correlate RELN genotype and secretion in humans we, thus, examined RELN levels in blood samples. Three RELN-immunoreactive bands with molecular masses of ∼430 (FL), 330 (NR6) and 160 (NR2) kDa were revealed (Fig. 4a, lanes 2-4) in blood samples of healthy individuals. The NR6 band was the most predominant form, whereas the smaller band NR2 was the less abundant (Fig. 4a) as previously reported for adult human, rat and mouse sera (*48*). Both the unprocessed RELN-FL protein and the two NR6 and NR2 proteolytic fragments co-migrated with recombinant RELN transfected into HEK293T cells (Fig. 4a, lane 1). The amount of all three RELN fragments were remarkably lower in the serum from the C2 patient than in that of the healthy mother (Fig. 4b). As missense mutations do not appear to affect protein synthesis, the lower levels of serum RELN most likely result from decreased secretion of the altered RELN proteins from the liver(*41*).

**Fig. 4.**
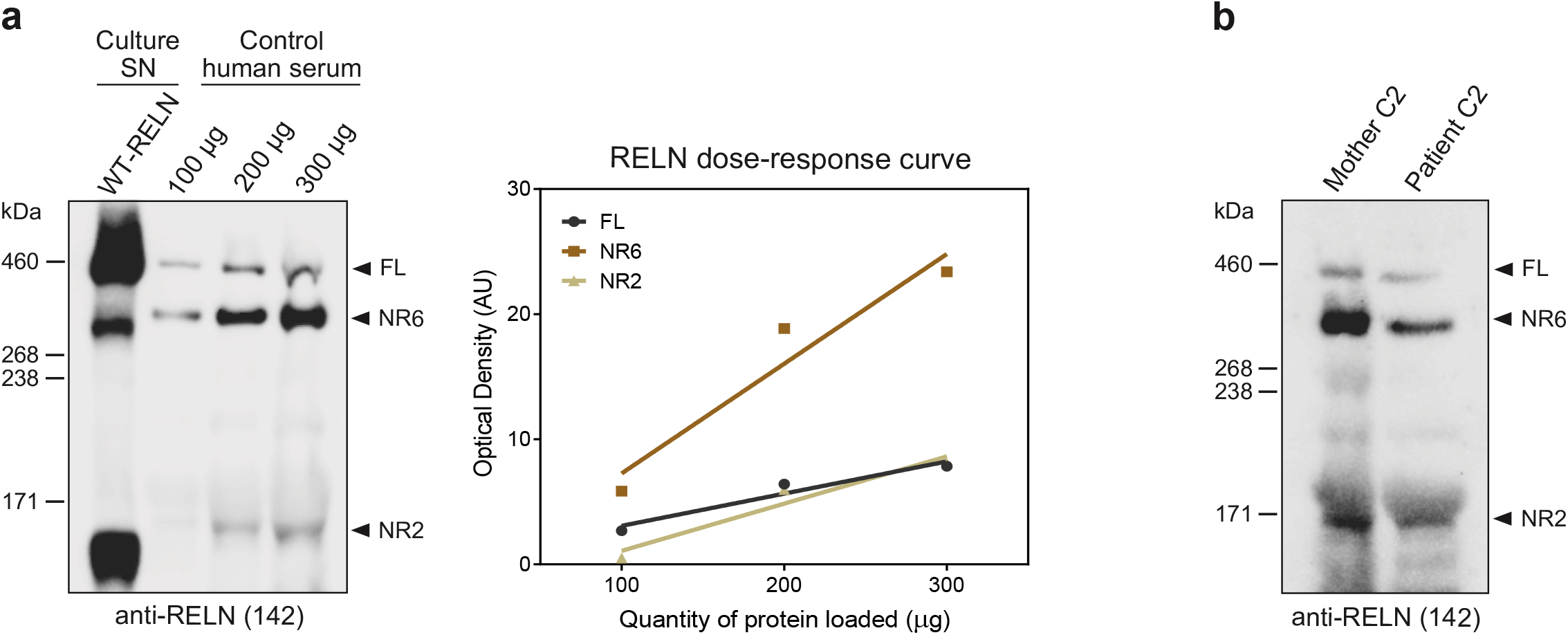
RELN levels are reduced in the blood serum from compound heterozygous patient C2 carrying I650S and D556V mutations. **a** Representative immunoblotting of blood serum RELN from control subjects (left panel) probed with anti-RELN 142 antibody with increasing protein quantities (μg). Recombinant WT-RELN from the supernatant (SN) of transfected HEK293T cells was loaded for reference. The right panel shows the optical densities (AU: arbitrary units) for all three RELN bands increasing linearly with the amounts of serum protein loaded. **b** Immunoblotting of blood serum RELN from the compound heterozygous patient C2 and the healthy mother showing a reduction in the amount of RELN in the patient. Protein standard sizes (kDa) are indicated on the left side of the blots.

Thus, the levels of circulating RELN confirm our previous *in vitro* findings whereby C2 mutations altered secretion and processing of RELN.

### *RELN* mutations affect the capacity to form aggregates along the antero-posterior axis of the developing cerebral cortex

In order to test whether *RELN* mutations affect its activity compared to their WT counterpart *in vivo* we took advantage of a functional assay developed by Kubo et al., (*51*). Ectopic RELN overexpression in the developing cortex of mouse embryos drives the formation of neuronal aggregates. These electroporated cells project their processes toward a central RELN-rich region poor in cell-bodies. Later-born neurons migrate through early-born neurons to reach the most internal part of this structure, recapitulating, even if ectopically, the inside-out development of the cortex. We electroporated plasmids carrying WT-RELN or the different point mutations followed by an IRES-eGFP in the developing cortex at E14.5 and collected the brains at P1 (Fig. 5a). As previously shown (*51*), we confirmed that WT-RELN is capable to induce the formation of aggregates (Fig. 5b). In addition, we found that these were not forming randomly along the rostro-caudal axis, but exclusively in caudal regions at hippocampal levels (n=5 for WT) (Fig. 5c). Different effects were obtained when mutations were electroporated. Y1821H, I650S and D556V identified in compound heterozygous and MI1/2 patients behaved as the WT with GFP^+^ aggregates forming caudally, while the G1280E and R913C mutations promoted the induction of aggregates at both caudal and rostral levels (Fig. 5b,c). Interestingly, the C539R and R3207C mutations found in patients DN1 and DN2 failed to form cell aggregates, consistent with their impaired secretion. For all mutations, aggregates were forming in the intermediate zone (IZ) just below the cortical plate (CP) labeled by Tbr1, a marker of deep-layer neurons at this age (Supplementary Fig. 3).

**Fig. 5.**
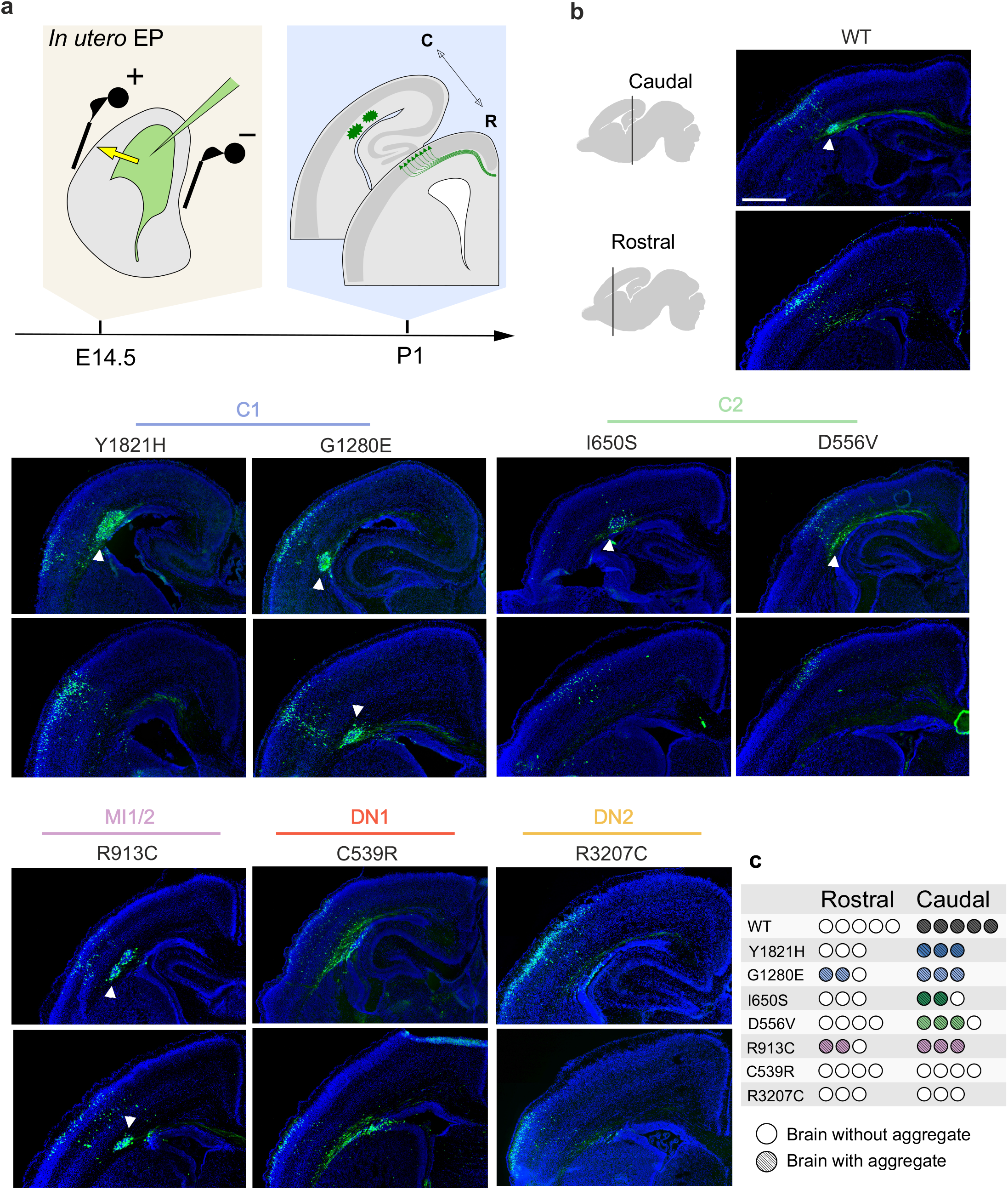
*RELN* mutations affect the capacity to form aggregates along the antero-posterior axis in the embryonic mouse cortex. **a** Schematic representation of *in utero* electroporation (IUE) at E14.5 and collection of the samples at P1. **b** Representative wide-field immunofluorescence images of GFP^+^ aggregates (DAPI counterstaining for nuclei in blue) at two rostro-caudal levels (Bregma 0.86 and -1.58) of P1 mouse brains upon IUE of WT-RELN and patients’ mutations. WT-RELN overexpression leads to the formation of aggregates only at caudal levels (n=5). Mutations belonging to the compound heterozygous C1: Y1821H behaves as the WT for aggregate distribution, while G1280E drives to the formation of aggregates also at rostral levels (n=3). Mutations belonging to C2: both I650S and D556V induce the generation of aggregates only at caudal levels (n=3 and n=4 respectively) similarly to the WT. The maternal-inherited mutation R913C of the MI1/2 patients drives the formation of aggregates at both caudal and rostral levels (n=3). *De novo* mutations C539R and R3207C of patients DN1 and DN2, respectively, fail to induce formation of aggregates either at caudal or rostral levels (n=4 and n=3 respectively). GFP^+^ neurons (green), DAPI staining (blue), white arrows indicate aggregates. **c** Quantification of aggregate formation for all electroporated constructs. Full circle represents a brain with aggregates, empty circle a brain without aggregates. Scale bar: 500 µm.

Overall, these results allowed to conclude that: i) aggregates are mostly obtained in posterior regions, indicating that different areas of the developing cortex are not equally responsive to RELN; ii) mutations of compound heterozygous patients lead to at least the formation of aggregates in the posterior cortex, indicating that they retain some of the activity of the WT protein; iii) mutations of C1 and MI1/2 patients appear to gain the capacity to induce aggregate formation at rostral levels and thus represent a GOF in this assay; iv) the two heterozygous *de novo* mutations behave as LOF as they completely abolish the ability to promote aggregate formation.

### Most of *RELN* missense mutations fail to form properly organized rosettes

When WT-RELN was electroporated the aggregates were organized recapitulating the inside-out formation of the cerebral cortex (*51*) and will be hence referred to as rosettes. In addition, about 50% of the rosettes (8/19) displayed a center that was cell-body-poor and accumulating RELN with the GFP^+^ electroporated cells projecting their processes towards it, analogously to the marginal zone (MZ)/layer I (LI) of the developing cortex (Fig. 6a,b and Supplementary Fig. 4). We thus asked whether the different missense mutations driving the formation of aggregates were actually able to generate properly formed rosettes. G1280E was the only mutation generating rosettes with a cell-body-poor central region 60% of the times (21/33) (Fig. 6a,b and Supplementary Fig. 4). Mutations Y1821H, I650S and R913C, instead, generated aggregates that could be considered rosettes-like structure as the GFP^+^ cells had their processes correctly directed towards a center in which there was an accumulation of RELN, but this center was invaded by late-born neurons expressing Brn2 (Fig. 6a,b and Supplementary Fig. 4). Moreover, the missense mutation D556V drove the formation of structures in which the GFP^+^ electroporated cells were spread throughout, without a proper organization and, although expressing RELN, the protein failed to accumulate it in a central region (Fig. 6a,b). This resulted in a structure completely lacking organization and a center, which we defined as an aggregate. Cells electroporated with the C539R and R3207C *de novo* mutations expressed RELN although they were unable to drive the formation of any sort of aggregates and some GFP^+^ cells appeared arrested in the VZ (Fig. 6a,b and Supplementary Fig. 4). Some of these neurons exhibited abnormally high levels of RELN intracellularly (Fig. 6, white arrows), confirming the impairment of secretion detected *in vitro*.

**Fig. 6.**
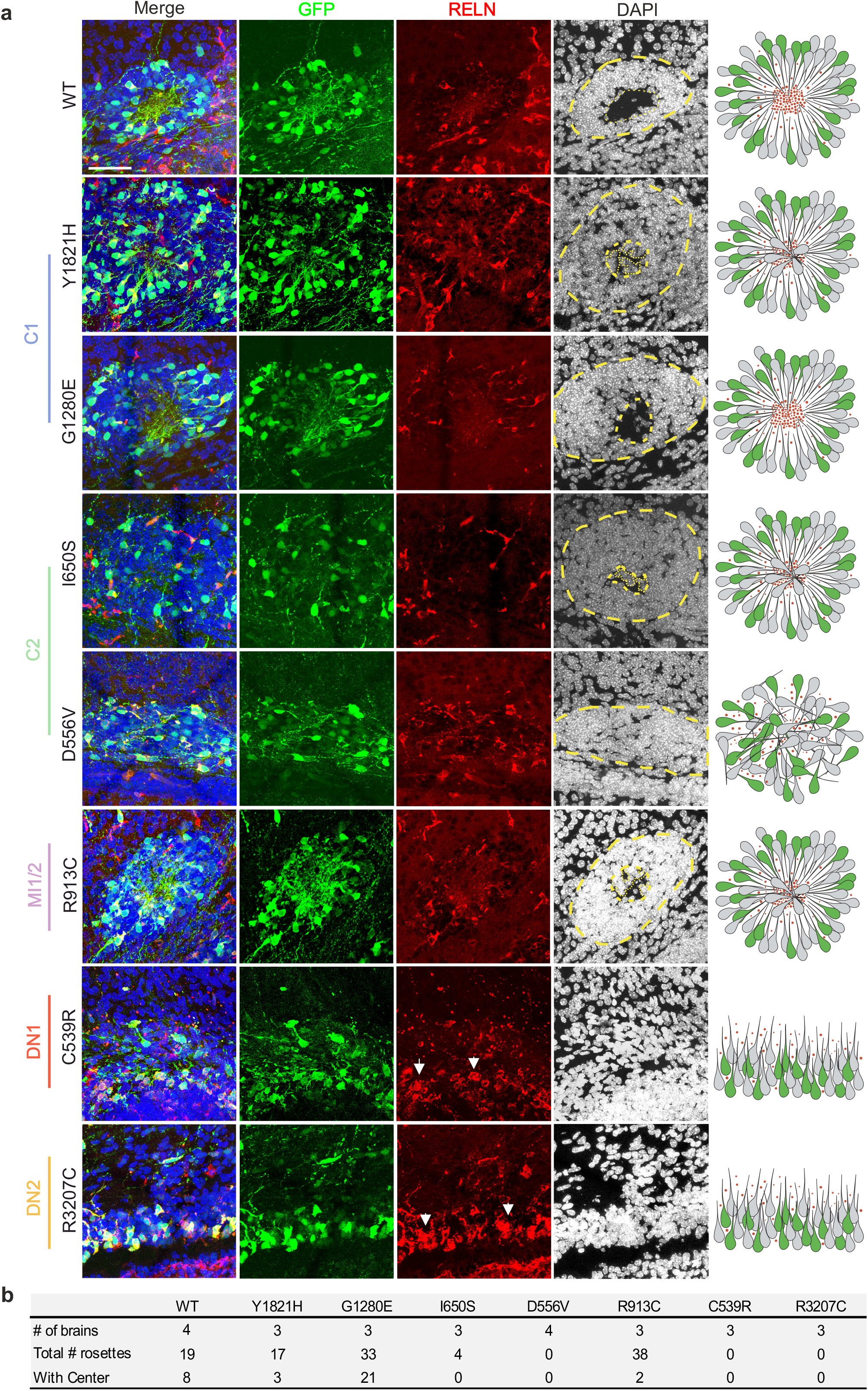
The majority of mutations fail to generate properly formed rosettes. **a** Mosaic maximum projection confocal images of aggregates. Aggregates with GFP^+^ electroporated cells projecting their processes toward a central region that is cell-body poor and RELN-rich are considered properly formed rosettes. Aggregates with GFP^+^ cells projecting their dendrites towards a central RELN-rich region but invaded by GFP^-^cells are considered rosettes with no center. Electroporation of WT-RELN or the G1280E mutation drive the formation of proper rosettes. Mutations Y1821H, I650S and R913C lead to the generation of rosettes with no cell-poor center. Mutation D556V fails to organize neurons correctly and leads to the formation of aggregates only. Mutations C539R and R3207C completely fails to form any sort of aggregate and they seem to accumulate more RELN in the cells (white arrows). Scale bar: 50 µm. **b** Quantification of the number of rosettes with a proper center and the total number of rosettes for WT-RELN and mutations. WT-RELN and mutation G1280E drive the formation of rosettes with a proper center in 42% and 63% of cases, respectively, while mutations Y1821H, I650S and R913C display a decreased capacity to form proper rosettes (18%, 0% and 5% respectively).

We conclude that all mutations, except for the G1280E mutation of C1, behave as LOF by altering the capacity to form rosettes or even aggregates with progressively more severe degrees, ranging from lack of cell-poor-centers (Y1821H, I650S and R913C mutations in patients C1, C2 and MI1/2 respectively) to lack of organization (D556V in C2) and complete loss of neuronal aggregation capacity (both *de novo* heterozygous mutations in DN1 and DN2 patients).

### *RELN* mutations alter neuronal migration rostrally

RELN is important to regulate neuronal migration (*2, 3*) and at rostral levels where rosettes are not normally forming, in the presence of WT-RELN electroporated GFP^+^ cells migrate to colonize the upper layers, in particular LII/III, accordingly to the stage of electroporation (E14.5) (Fig.7 left panel). We divided the CP in 10 equal bins and quantified the percentage of GFP^+^ cells per bin. Bin 1 corresponded to the MZ/LI, bin 2-4 to the upper layers (UL), bin 5-7 to the deep layers (DL), bin 8-9 to the IZ and bin 10 to the VZ. When WT RELN was overexpressed, 90% of GFP^+^ pyramidal neurons were found within bin 2-3 (70% in bin 2 and 20% in bin 3), corresponding approximately to LII/III as expected by the stage of electroporation. The remaining 10% of GFP^+^ cells were spread in the other bins (Fig. 7a left panel and 7b). When the Y1821H and I650S mutations were tested, a delay was observed in the migration of the electroporated cells, with significantly less cells in bin 2 and more in bin 3. The G1280E and R913C mutations, instead, promoted an increase in the number of GFP^+^ cells trailing specifically in bin 6 and 7/10, respectively, corresponding to DL, despite no significant decrease in the number of neurons able to reach the UL (Fig. 7a,b). D556V and R3207C mutations did not affect the migration of the electroporated cells, thus behaving as the WT-RELN. The most striking effect was observed for the C539R mutation, which severely delayed electroporated GFP^+^ cells with only 50% of them reaching the UL (bins 2 and 3) and the remaining detected in deep locations, in particular in bins 7 to 10 (Fig. 7a,b), corresponding to DL (layer V/VI), IZ and VZ.

**Fig. 7.**
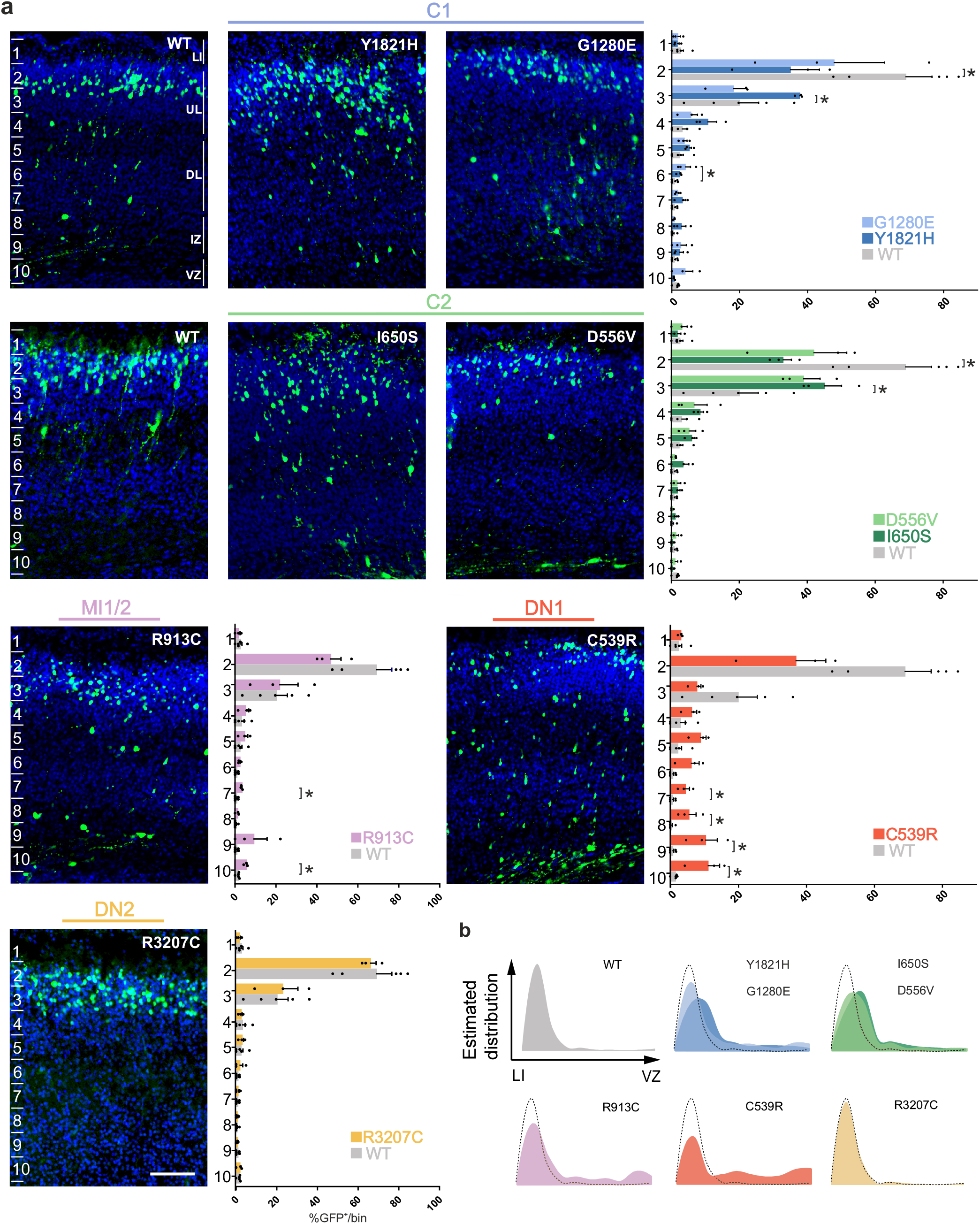
*RELN* mutations affect cell migration at rostral levels. **a** Immunofluorescence wide-field images of P1 brains after IUE at E14.5. A portion of the electroporated cortex was divided into 10 bins and the percentage of electroporated GFP^+^ cells per bin was calculated. Bin 1 corresponded to LI, bin 2-4 to the upper layers (UL), bin 5-7 to the deep layers (DL), bin 8-9 to the intermediate zone (IZ) and bin 10 to the ventricular zone (VZ). As expected by the stage of electroporation, GFP^+^ neurons migrate to colonize the UL when WT plasmid is overexpressed with approximately 90% of GFP^+^ neurons found in bin 2 and 3 (gray graphs). When mutation Y1821H is overexpressed, electroporated neurons are delayed in their migration with significantly more GFP^+^ cells found in bin 3 and less in bin 2 (*p*=0.035 and 0.035 respectively). For mutation G1280E there is an increase in GFP^+^ cells only in bin 6, while the distribution in the other bins is similar to that of the WT. Mutation I650S drives a delay of electroporated neurons with more neurons in bin 3 and less in bin 2 compared to the WT (*p*=0.035 and 0.035 respectively), while mutation D556V do not affect migration of GFP^+^ cells. Mutation R913C induces a delay in migration with more cells in bin 7 and 10 (*p*=0.035 and 0.035 respectively). Mutation C539R severely affects the migration of electroporated neurons, with an increase in the number of cells found in bin 7 to 10 (*p*=0.035, 0.017, 0.035 and 0.035 respectively). Mutation R3207C does not induce any delay in the migration of electroporated cells. **b** Recapitulative representation of the estimated distribution of electroporated cells from the marginal zone (MZ) to the ventricular zone (VZ). Scale bar: 100 µm.

In order to study whether delayed migration was accompanied by changes in morphological features or fate we analyzed both the cells that were delayed in the CP and those able to reach the correct position in the UL. Delayed cells for all mutations displayed a morphology of migrating neurons with a long apical process accumulating RELN (Supplementary Fig. 5a). D556V, C539R and R3207C mutations appeared to induce an increased accumulation of RELN inside the cytoplasm of GFP^+^ cells (Supplementary Fig. 5a white arrows) correlating with the *in vitro* observations (Fig. 2b). Those that were able to reach the UL for both WT-RELN and the different mutations appeared to differentiate normally into pyramidal neurons having their dendrites in layer I and accumulating RELN mainly in the primary apical dendrite (Supplementary Fig. 6a). Both delayed GFP^+^ cells in the CP and those arrived in the upper CP maintained the identity of Brn2^+^ upper layer neurons for every mutation as for overexpression of the WT RELN (Supplementary Fig. 5b,6b) showing that although mispositioned electroporated cells maintained the correct fate.

We conclude that a majority of mutations alter the migration of electroporated cells at rostral levels, although to different degrees with the *de novo* C539R mutation of DN1 being the most severely impaired. Notably, the DN2 R3207C mutation, which behaved similarly to C539R in all other *in vitro* and *in vivo* assays, did not perturb neuronal migration.

Altogether, these results show that *RELN* missense variants alter different aspects of RELN processing and function (Table 1). In particular, defects of *in vitro* secretion and *in vivo* regulation of neuronal aggregation and/or migration correlate with the patients’ phenotype and provide molecular insights on the cause of a broad spectrum of NMDs.

**Table 1:**
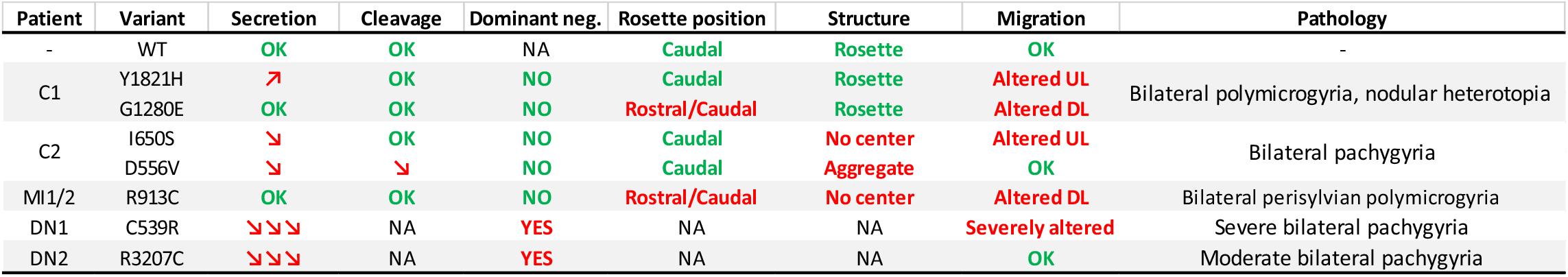
Summary of performed assays. *In vitro* and *in vivo* assay results and correlation with patients’ phenotypes. UL: upper layers, DL: deep layers.

| Patient | Variant | Secretion | Cleavage | Dominant neg. | Rosette position | Structure | Migration | Pathology |
| --- | --- | --- | --- | --- | --- | --- | --- | --- |
| - | WT | OK | OK | NA | Caudal | Rosette | OK | - |
| C1 | Y1821H | ↗ | OK | NO | Caudal | Rosette | Altered UL | Bilateral polymicrogyria, nodular heterotopia |
|  | G1280E | OK | OK | NO | Rostral/Caudal | Rosette | Altered DL |  |
| C2 | I650S | ↘ | OK | NO | Caudal | No center | Altered UL | Bilateral pachygyria |
|  | D556V | ↘ | ↘ | NO | Caudal | Aggregate | OK |  |
| MI1/2 | R913C | OK | OK | NO | Rostral/Caudal | No center | Altered DL | Bilateral perisylvian polymicrogyria |
| DN1 | C539R | ↘↘↘ | NA | YES | NA | NA | Severely altered | Severe bilateral pachygyria |
| DN2 | R3207C | ↘↘↘ | NA | YES | NA | NA | OK | Moderate bilateral pachygyria |

## DISCUSSION

Until now, *RELN* mutations have been associated to a wide spectrum of neurodevelopmental disorders ranging from recessive forms of NMDs, namely lissencephaly with cerebellar hypoplasia (LCH) (*52*), to dominant form of epilepsy (ADTLE) (*41*), or psychiatric disorders such as autism and schizophrenia (*38*). RELN is a pleiotropic factor, now being shown to be a key regulator of multiple biological processes such as neuronal migration, the formation of layered structures and synaptogenesis. These recent observations suggest that the spectrum of RELN related disorders might reflect the degree of severity of *RELN* mutations on protein function. However, the high mutation rate due to the large size of the *RELN* gene and the extensive variety of associated phenotypes contributed to question these variants as significant in the etiology of these disorders. Hence, the relevance of heterozygous mutations in the function of the RELN protein underlying these distinct pathological conditions remained unexplored. Here, we show that missense *RELN* mutations are deleterious for the function of the RELN protein and can be associated with a broad spectrum of recessive and dominant NMDs. Indeed, using complementary *in vitro* and *in vivo* assays we were able to correlate specific misfunctions of the RELN protein, both LOF and GOF, and their severity with the patients’ phenotypes.

We report here 6 patients with a spectrum of neurodevelopmental disorders ranging from severe polymicrogyria with nodular heterotopia to milder pachygyria. None of these patients exhibited any of the cerebellar anomalies previously considered as a hallmark of the *RELN* associated autosomal recessive lissencephaly with cerebellar hypoplasia (LCH) (*52*). Our data associate for the first time *de novo* monoallelic *RELN* missense mutations with cortical malformations. So far, heterozygous *RELN* mutations had only been associated with psychiatric disorders, ASD or ADTLE (*38, 41*) and only one mutation has been functionally tested (*42*). Indeed the heterozygous *reeler* mouse mutant with 50% reduction of the protein is considered a model for schizophrenia (*31*) and results in altered cortical circuits, but normal layering, possibly explaining the insurgence of ADTLE or psychiatric disorders in humans. Less obvious is how heterozygous mutations can lead to disorganization of cortical layering, as observed in patients of this study, a process that is not affected even with a 50% reduction of RELN (*31*), suggesting that the type of mutation can correlate with the inheritance and severity of the phenotype. Indeed, in the majority of reported patients with NMDs (*33-35*), *RELN* mutations caused protein truncation or a null allele. In our patients, they are missense with a full-length protein generated, but not properly functioning.

Among the 6 MCDs patients, while three (C2, DN1 and DN2) share a similar phenotype consisting of frontally predominant pachygyria, two (C1 and MI1/2) show different phenotypes, featuring a combination of polymicrogyria with extensive bilateral periventricular nodular heterotopia in one and bilateral perisylvian polymicrogyria in the other. These differences do not correlate with the transmission pattern of the mutation, biallelic or monoallelic, but rather reflect the impact of the effect of such mutations on RELN function and, thus, on the developing cortex. According to the developmental and genetic classification for MCDs (PMID: 22427329 (*53*)), pachygyria belongs to the group of ‘Malformations Secondary to Abnormal Neuronal Migration’, while polymicrogyria is included in ‘Malformations Secondary to Abnormal Postmigrational Development’, although it is recognized that distinction between these two malformations is not always feasible using MRI and that these entities might be combined. Nodular heterotopia that is instead supposed to result from abnormalities of the initiation of migration (*54*). Indeed, neuronal migration involves several steps during development, which are all RELN-dependent. The newly generated neurons have to attach to the radial glia that will guide their first type of migration called locomotion. When neurons reach the correct place in the cortical plate they have to detach from the radial glia and complete their migration with a terminal somal translocation (*55, 56*). Based on this hypothesis, the malformation spectrum we observed could reflect the pleomorphic effects of *RELN* mutations affecting, according to their cellular consequences, neuronal migration either at its initial stages or later during the transmantle migration. We performed a series of assays to test different aspects of RELN function, in particular its capability to form neuronal aggregates, its effect on cell migration as well as its secretion and cleavage in order to dissect how the different missense mutations could affect these functions, thus resulting in the different malformations observed in the patients. Our results indicate that the polymicrogyric cortex and nodular heterotopia could originate from defects in the attachment to and/or disengagement from radial glia at different time points, resulting in an inability of neurons to enter or locate properly in the cortical plate (*57*). Pachygyria would result from impairment in key steps of migration (initiation, locomotion, somal translocation) with a severity scaled to the accumulation of multiple affected stages.

We have shown that the 4 mutations belonging to the compound heterozygous patients alter at some level RELN ability to promote aggregates formation even if with different degrees. The Y1821H mutation in patients C1 and the I650S in patient C2 correctly drive the formation of rosettes only in caudal regions, however they lack the capacity to stop the later-born neurons to invade the central region. In addition, both mutations also alter the migration of electroporated cells at rostral levels, indicating that they are indeed affecting the RELN capacity to cell autonomously regulate neuronal migration. The second mutation instead behaves differently in either patient. The G1280E mutation detected in patient C1 drives the formation of properly formed rosettes, but both in caudal and rostral regions behaving as a GOF, and it delays neuronal migration in deep layers. The D556V mutation of patient C2 promotes the generation of aggregates that lack the capacity to organize the electroporated cells, while it does not influence neuronal migration. Moreover, the G1280E variant is present in around 1% of the normal population, with 26 reported homozygous individuals in the gnomAD database (*58*), pointing to the fact that alone it is not sufficient to drive a pathological condition. Thus, it is the combined effect of the two mutations of each compound heterozygous patient that might contribute to the observed phenotypes. This is further supported by the consequences on secretion and processing of the mutated proteins since mutations of the two compound heterozygous patients show opposite behaviors, with one mutation of the C1 patient exhibiting an increased amount of secreted RELN (Y1821H) and both mutations of patient C2 decreasing it. In particular, the D556V mutation, although distant from the two main cleavage sites (N-t cleavage occurs between P1244 and A1245, while C-t cleavage between R3455 and S3456) (*6, 7, 59*), specifically reduces the C-t cleavage of the protein. The substitution of a negatively charged amino acid with a neutral one could have an impact on the folding of RELN, thus preventing the protein to acquire the proper conformation to be correctly cleaved or to bind to its receptors (VLDLR and ApoER2). A recent study (*60*) described a similar phenotype after overexpression of RELN in a Nrp1 knockdown (KD) mouse model. The authors demonstrated that Nrp1 forms a complex with VLDLR to which RELN strongly binds and its knocked-down results in a misorientation of electroporated cells in the aggregates. The misfolded protein carrying the D556V mutation could be less efficient in binding to the receptors, explaining why when overexpressed *in vivo*, although retaining the capacity to form aggregates, it loses the ability to correctly organize them. Notably, cotransfection of I650S and D556V also resulted in decreased RELN secretion and correlated with the reduction observed in the serum of the C2 patient, suggesting a LOF effect caused by impaired secretion of the altered proteins. Overall, the compound heterozygous patients exhibit a less severe phenotype compared with the classical LCH lacking the protein (*33*), as the 4 mutant proteins are still present in the secreted fraction and partially retain the capacity to form aggregates.

The R913C mutation observed in patients MI1/2 is maternally inherited with the mother having had epilepsy during childhood. This mutation has a dominant pattern of inheritance as previously described in patients with ADTLE (*41*). Nevertheless, since whole genome sequencing has yet to be performed, other factors cannot be ruled out. This mutation promotes the formation of rosettes also in rostral region like the G1280E mutation, however is not capable to retain the cells to invade the central RELN-rich region similarly to Y1821H and I650S mutations. The R913C mutation delays neuronal migration in deep layers, again similarly to G1280E. Thus, it affects both RELN capability of regulating neuronal migration and responsiveness to permissive cues along the rostro-caudal axis. Interestingly, both M1/2 and C1 patients display polymicrogyria and the R913C and G1280E mutations they harbor behave alike in *in vivo* assays.

The two *de novo* heterozygous patients DN1 and DN2 exhibit phenotypes of different severity. Both the C539R and R3207C mutations when electroporated in the developing mouse cortex result in complete LOF, abrogating the formation of organized rosettes or even aggregates. Our results also show that C539R and R3207C can act as dominant negative forms, impairing not only their own secretion, but also that of the protein produced by the WT allele. However, exclusively the overexpression of C539R severely affects the migration of the electroporated cells rostrally, with the majority of cells delayed in the cortical plate or arrested in the VZ correlating with the severity of the phenotype. The C539R mutation is found in the RELN N-t region that was shown to interact with α3β1 integrin regulating neuronal migration (*61*). Aberrant folding of this region could impair this interaction thus determining the neuronal migration defect observed *in vivo*. The R3207C mutation is not localized in a region thought to be important for binding to any known RELN receptor, thus correlating with its normal behavior in the migration assay. Both mutations induce a change from a Cysteine (Cys) to an Arginine (Arg) or viceversa. Cys are not only important for homodimerization (*12*) but also for intramolecular disulfide bridges (*62*). The C539R mutation affects a Cys, shown to form an intramolecular disulfide bond with the Cys^463^, thus impairing the formation of this bridge and the RELN tertiary structure. Moreover, Cys^463^ is left free to form disulfide bonds and could interact with other Cys on the WT protein, thus explaining the dominant negative phenotype. The R3207C mutation is introducing a new Cys into the protein that could again affect the correct folding (*62*) and interact with the WT protein forming intermolecular bonding, thus reducing not only its own secretion, but also that of the WT protein. Although reports on RELN homodimerization have only focused on the extracellular space (*11*), we have demonstrated that it also occurs intracellularly. We, thus, propose that severely altering the tertiary conformation of RELN could drive to an abnormal intracellular dimerization, correlating with the observed accumulation of the protein inside the cells and, consequently, the dominant negative effect.

Affecting a crucial Cys for proper folding, could also contribute to the most severe phenotype observed in patient DN1. However, the difference in phenotypes observed in patients DN2 and MI1/2, with the R3207C mutation (DN2) associated with pachygyria and behaving as a dominant negative form, and the R913C (MI1/2) with polymicrogyria, pinpoints the fact that it is not sufficient to introduce randomly a Cys to result in a complete LOF of the protein, but some positions are more critical than others.

In conclusion, we have for the first time functionally characterized both *in vitro* and *in vivo* the effects of novel *RELN* mutations and correlated the severity of patients’ phenotypes with the levels of alteration of the RELN protein. Moreover, RELN serum levels in patient C2 support our *in vitro* findings highlighting the relevance of circulating RELN for diagnosis. Lastly, our results demonstrate that *RELN* missense mutations cause cortical malformations not only with recessive but also dominant inheritance.

This study paves the road to functionally analyze future identified *RELN* mutations to determine their involvement in such pathologies for genotype-phenotype diagnostics. Since RELN perfusion in the brain has been shown to recover loss-of-function behavioral defects in the mouse(*63, 64*), this work could open the possibility for intervention by allowing novel strategies for therapy.

## MATERIALS AND METHODS

### Subjects

A total of 6 pediatric patients that presented radiologic evidence of abnormal cortical development were investigated. This is a selected cohort comprising patients who were ascertained directly by the authors (NBB, CB, RG, EP, DJ) or whose clinical information and brain magnetic resonance images (MRIs) were sent to the authors. Brain MRI were reevaluated by an expert neuroradiologist trained in cortical malformations (CJR). Genomic DNA was extracted from blood samples. Venous blood (5-10 mL) was drawn into BD Vacutainer® SST™ tubes and then centrifuged at 1300 *g* for 10 min at RT. Plasma was collected, aliquoted and frozen at -80°C until assay. Genetic testing was performed in trios using a custom next-generation sequencing (NGS) panel targeting genes associated with MCDs (list of genes available upon request). The consanguineous family of MI1 and MI2 was also sequenced by whole-exome sequencing (WES) using Agilent SureSelectXT Clinical Research Exome (SureSelectXT Human All Exon V5 baited with clinically relevant genes) and the Illumina NextSeq 550 sequencer (Illumina, Inc., San Diego, CA, USA) according to the manufacturer’s protocol.

### Animals

Pregnant SWISS mice were purchased from Janvier Lab. All animals were handled in strict accordance with good animal practice as defined by the national animal welfare bodies, and all mouse work was approved by the French Ministry of Higher Education, Research and Innovation as well as the Animal Experimentation Ethical Committee of Paris Descartes University (CEEA-34, licence numbers: 18011-2018012612027541 and 19319-2018020717269338).

### Construction of plasmids

The full-length mouse *RELN* expression construct pCrl was kindly provided by Drs T. Curran and F. Tissir. The *RELN* cDNA was inserted using EcoRI and NotI restriction sites into a pCAG-IRES-NLS-EGFP vector containing a modified chicken β-actin promoter and a cytomegalovirus-immediate early enhancer (CAG)(*65*). Point mutations were inserted using either the In-Fusion® HD Cloning Kit (Takara) according to the manufacturer protocols directly into the pCAG vector, or by PCR in pCrl, and then subcloned in the pCAG vector. Plasmid amplification was achieved using an EndoFree Plasmid Maxi Kit (Qiagen) and due to the large size (16 kb) special care was taken to avoid mechanical damage during the procedure. The mouse protein is 94.2% identical to the human protein at the amino acid (aa) level and it has 3461 amino acids *versus* the 3460 in humans with an additional aa at the N-terminus. The concerned missense mutations were all found conserved in the mouse sequence but at position +1aa. For this reason, the different human mutations were cloned in position +1aa into the mouse *RELN* cDNA. The obtained plasmids were verified by restriction and sequencing to confirm that the right point mutation was correctly inserted.

### Cell culture, transfection and sample preparation

Human embryonic kidney (HEK) 293T cells were grown in Dulbecco’s modified Eagle’s medium (DMEM; Gibco) containing 10% fetal bovine serum (FBS) and 1% penicillin-streptomycin (Gibco) and maintained at 37°C in a 5% CO_2_ atmosphere. Cells were seeded onto 6-multiwell plates at a density of 62500 cells/cm^2^. Twenty-four hours later, transfection with 5 μg pCAG constructs containing wild-type (WT) *RELN* or different *RELN* mutants (Y1821H, G1280E, I650S, D556V, R913C, C539R, R3207C) was performed for 4 hrs using Lipofectamine 2000 (Invitrogen) according to the manufacturer’s instructions. Mock-transfected cells (treated with transfection reagent only) and cells transfected with a pCAG-GFP plasmid were used as controls. To phenocopy the heterozygous patients’ genotypes, cells were co-transfected with 2.5 μg of each pCAG construct. Cells were maintained in serum-free Opti-MEM (Gibco) supplemented with 1% antibiotics after transfection and cultured for an additional 40 hrs. Conditioned media were collected and a protease inhibitor cocktail (cOmplete™ tablets, Roche, Germany) was immediately added. After clearing by centrifugation at 10000 rpm for 10 min, samples were concentrated using Amicon Ultra 50K centrifugal filters (Millipore) at 4000 *g* for 30 min and stored at -80°C. Cell lysates from transfected cells were collected using RIPA buffer (50 mM Tris-HCl pH 7.5, 150 mM NaCl, 1% NP-40, 0.1% SDS, 0.5% sodium deoxycholate) supplemented with 2 mM EDTA and protease inhibitors (cOmplete™ EDTA-free tablets). Samples were then left in a rotator for 30 min at 4°C to better dissolve the proteins and centrifuged at max speed for 10 min at 4°C in order to pellet out crude undissolved fractions. The supernatant was stored at -20°C until use. Protein quantification was determined using the bicinchoninic acid (BCA) protein assay reagent kit (Pierce™, ThermoFisher Scientific, USA) using bovine serum albumin (BSA) as standard.

### Western blot analysis

Proteins samples were added to 1/4 volume of 4X LDS sample buffer (141 mM Tris, 106 mM Tris-HCl, 2% LDS, 10% glycerol, 0.51 mM EDTA, 0.22 mM SERVA Blue G, 0.175 mM phenol red, pH 8.5), to 1/10 of sample reducing agent 10X (50 mM DTT) and boiled at 95°C for 3 min. For non–reducing conditions no DTT was added. Protein samples (10 µg of cellular fraction; 1 µg of secreted fraction; 200 µg of human serum) were separated by SDS-PAGE on 3-8% tris-acetate gels (NuPAGE, Invitrogen) under reducing conditions at 150 V (cellular fraction) or 90 V (secreted fraction and serum) at room temperature (RT) and electro-transferred to 0.45 µm nitrocellulose membranes (Amersham, GE Healthcare, Germany) for 1h30-2h at 0.5 A at 4°C. After 1 h blocking at RT with 5% (w/v) milk or BSA in Tris-buffered saline (50 mM Tris, 150 mM NaCl, pH 7.6) containing 0.1% Tween 20 (TBS-T), membranes were incubated overnight at 4°C with the following primary antibodies: mouse anti-RELN G10 (MAB5364, Millipore 1:2000) or 142 (for human RELN, MAB5366, Millipore 1:400), directed against the N-terminal region, a mix of mouse anti-RELN clones 12 and 14 (1:250, gift from Dr. André Goffinet) that recognize epitopes in the C-terminal(*44*), rabbit anti-GFP (A6455, Invitrogen 1:2000) and mouse anti-panCadherin (C1821, Sigma 1:2000) as loading controls (cellular fraction). After three washing periods of 10 min with TBS-T, the membranes were incubated with the appropriate HRP-conjugated secondary antibodies (Jackson Immunoresearch; 1:20000) diluted in TBS-T with 5% (w/v) milk for 1h at RT. After three 10 min washes with TBS-T, blots were developed with SuperSignal West Pico Chemiluminescent Substrate or with SuperSignal West Femto Maximum Sensitivity Substrate (ThermoScientific) and visualized on the ChemiDoc apparatus (Bio-Rad). Quantitative analysis was determined by densitometry using Image Lab™ software (Bio-Rad). The densities of protein bands were quantified with background subtraction. The bands of the cellular fraction were normalized to GFP loading control while those of the secreted fraction were normalized to total protein (Ponceau S). The molecular weights were determined by using an appropriate pre-stained protein standard (HiMark 31-460 kDa, Invitrogen) for high molecular weight proteins.

### *In utero* electroporation

E14.5 timed-pregnant Swiss mice were subjected to abdominal incision to expose the uterine horns under anesthesia with Isoflurane (AXIENCE SAS) at a concentration of 4% for induction and 2% for maintenance. A subcutaneous injection of buprenorphine (0.05 µg/g) and ketoprophen (5 µg/g) was delivered as pre- and post-op analgesia, respectively. Plasmids carrying the WT *RELN*, or the different mutations were injected at a concentration between 4 and 7 µg /µl through the uterine wall into one of the lateral ventricles of each embryo by a glass pipet. After soaking the uterine horn with a PBS solution, the embryo’s head was carefully held#between a pair of electrodes. 5 electrical pulses of 40V and 50ms with an interval of 1s were delivered using a NEPA21 electroporator (Nepagene). The uterine horns were returned into the abdominal cavity after electroporation, and embryos were allowed to continue their normal development up to birth.

### Tissue preparation and immunohistochemistry

The birth date was considered as postnatal day 0 (P0). P1 animals were decapitated and the brains were dissected and fixed overnight by immersion in 4% paraformaldehyde (PFA) in 0.1M phosphate buffer (PB), pH 7.4 at 4°C. The next day brains were cryoprotected with 20% sucrose in PB solution overnight at 4°C. Brains were then embedded in OCT and sectioned in 16 µm slices. Immunostaining was performed as previously described (Bielle et al., 2005; Griveau et al., 2010; de Frutos et al., 2016). Primary antibodies used for immunohistochemistry were: mouse anti-RELN G10 (MAB5364, Millipore 1:1000), chicken anti-GFP (GFP-1020, Aves Labs, 1:2000), goat anti-Brn2 (sc-6029 Santa Cruz biotechnology, 1:500), rabbit anti-Tbr1 (ab31940, Abcam, 1:1000). Secondary antibodies used were: donkey Cy5 anti-goat (705-175-147, Jackson ImmunoResearch Laboratories, 1:500), donkey Cy3 anti-rabbit (711-165-152, Jackson ImmunoResearch Laboratories, 1:700), donkey Alexa-488 anti-chicken (703-545-155, Jackson ImmunoResearch Laboratories, 1:1000), donkey Alexa-555 anti-mouse (A-31570, Molecular Probes, 1:1000). DAPI (5 μg/ml, D1306, ThermoFisher Scientific) was used for fluorescent nuclear counterstaining of the tissue and mounting was done in Vectashield (H-1000, Vector Labs).

### Image acquisition and data treatment

Immunofluorescence images were acquired using a slide scanner Nanozoomer 2.0 (Hamamatsu) with a 20x objective or a Leica TSC SP8 inverted confocal microscope with 20x or 63x oil objectives (with or without digital zoom). Confocal images consist of multiple tile regions (mosaics) combined with serial z-stacks (0.5 µm through all the section’s depth). Composite images are presented as maximum projections and were generated by the LAS X software using Mosaic Merge and Projection functions.Discontinuities are due to mosaic stitching algorithm. For the migration assay, a portion of the electroporated section was divided in 10 bins and the number of GFP^+^ neurons, detected by immunofluorescence, in rostral regions for each condition was counted using Adobe Photoshop CS4 software in order to obtain the percentage of GFP^+^ cells per bin.

### Statistical analysis

Statistics and plotting were performed using GraphPad Prism 7.0 (GraphPad Software Inc., USA). Data are presented as mean ± SEM. Statistical comparison of the results of total RELN obtained by western blot analysis was performed using unpaired one sample Student’s *t* test (hypothetical value of 1) after they passed the Shapiro-Wilk normality test. Cleavage statistics were performed using nonparametric two-tailed Mann-Whitney *U* test. For the migration assay, the non-parametric Kolmogrov-Smirnov test to compare cumulative distributions was performed. *P* values < 0.05 were considered statistically significant and set as follows: \**p*<0.05, \*\**p*<0.01, \*\*\**p*<0.001.

## ACKNOWLEDGEMENTS

The authors wish to thank O. Gribouval for help with human genetics, N. Boddaert, AG. Lemoing, A. Toussaint and J.Steffann for recruitment of patients and diagnosis, T. Curran, the St. Jude Children’s Research Hospital (Philadelphia) and F. Tissir for the original mouse Reelin cDNA in pCrl, A. Goffinet for the RELN antibodies, W. Dobyns and N. Di Donato for their expertise in cortical malformations and helpful discussions at the onset of the project, the NeuroImag platform at the IPNP and SFR Necker imaging and histology platforms at the *Imagine* Institute for help with acquisition, the Animalliance platform for animal care. We are grateful to the patients and their families for their contribution to our research, P. Billuart and A. Cwetsch as well as members of the Pierani lab for technical support and helpful discussions, P. Bun from NeuroImag for help in image processing and C. Antignac, MC. Angulo and M. Cavazzana for critical reading of the manuscript.

## Funding

French Ministry of Research (BioSPc Doctoral school) (MR)

Fondation pour la recherche médicale, FDT20201201037 (MR)

Centre national de la recherche scientifique (CNRS) (AP)

Agence Nationale de la Recherche, ANR-15-CE16-0003-01 and ANR-19-CE16-0017-03 (AP)

Fondation pour la recherche médicale, Équipe FRM DEQ20130326521 and EQU201903007836) (AP)

Agence Nationale de la Recherche under “Investissements d’avenir” program, ANR-10-IAHU-01) (*Imagine* Institute=.

## Authors contribution

Conceptualization: MR, SF, FC and AP

Clinical assessment: RG, NBB, DJ, EP, EF, CB, CJR

Methodology: MR, SF, VM, OH, FC

Investigation: MR, SF and AP

Data Curation: MR, SF, NBB and AP

Writing – original draft preparation: MR, SF, NBB and AP

Writing – review and editing: MR, SF, FC, DJ, EP, RG, NBB and AP

Visualization: MR, SF, FC, NBB and AP

Supervision: AP

Project administration: AP

Funding acquisition: AP

