## Supplementary Data for "Functional characterization of RELN missense mutations involved in recessive and dominant forms of Neuronal Migration Disorders"

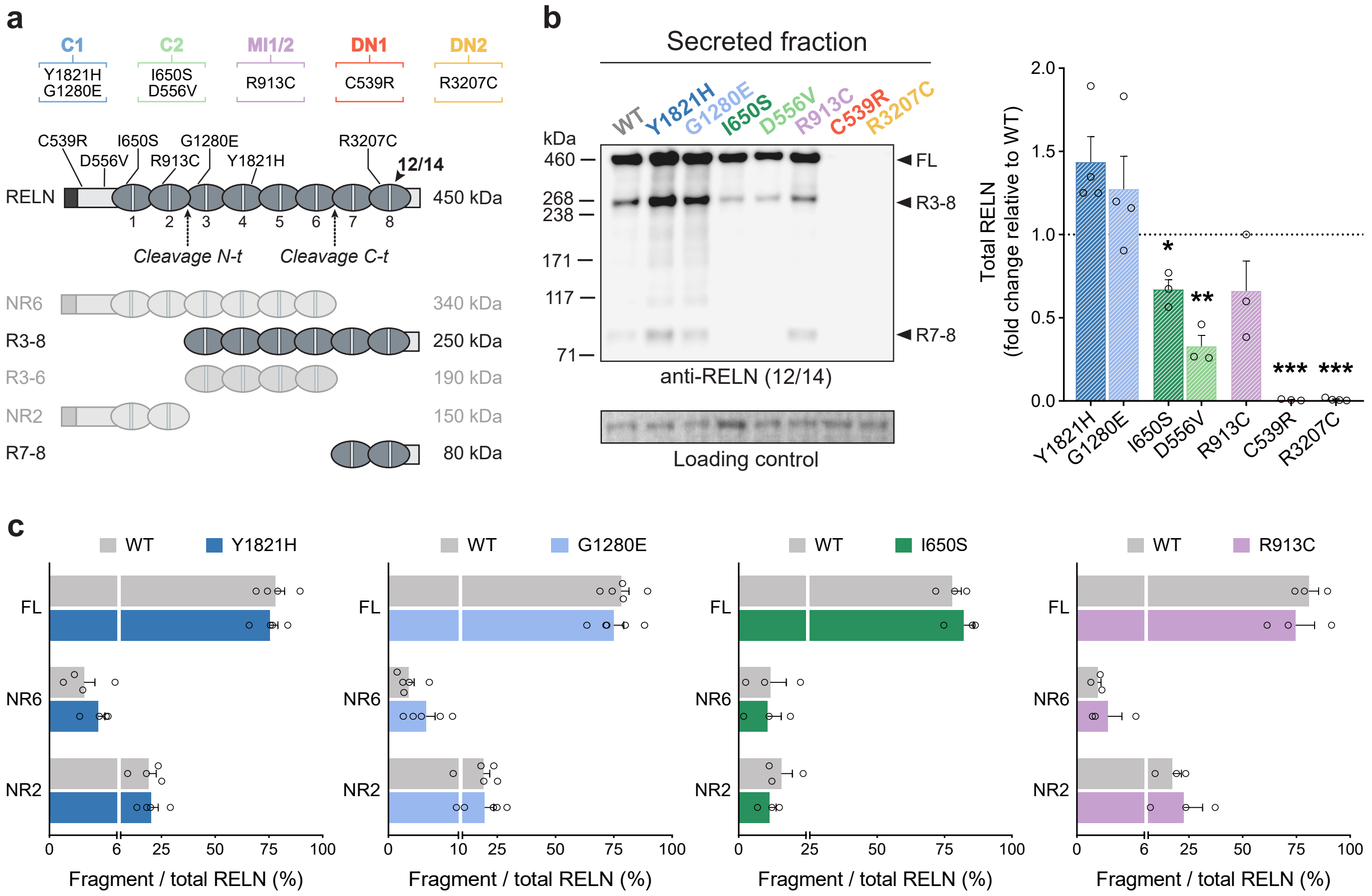


**Supplementary Fig. 1 Alterations of RELN secretion *in vitro* by missense mutations determined with antibodies 12/14 and protein cleavage.** **a** Schematic of the full-length RELN protein (450 kDa), its cleavage sites N-t and C-t (dotted arrows), and its five cleaved products (NR6, R3-8, R3-6, NR2, R7-8). The binding region of the 12/14 antibody and the position of *RELN* mutations are indicated with a black arrow and lines respectively. **b** Representative immunoblotting of the secreted fraction of HEK293T cells transfected with either RELN-WT or RELN-mutants, probed with anti-RELN 12/14 antibody. Right panel shows the densitometric analysis of total RELN levels normalized to total protein (Ponceau S) (n=3-4 independent experiments). Decreased levels of RELN were detected in media of cells transfected with mutants I650S (*$p$=0.0307), D556V (**$p$=0.0095), C539R (***$p$<0.0001) or R3207C (***$p$<0.0001). Protein standard sizes (kDa) are indicated on the left side of the blots. **c** Densitometric analysis of the proportion (%) of RELN-FL, NR6 (from C-t cleavage) and NR2 (from N-t cleavage) relative to the total amount of secreted Y1821H, G1280E, I650S, R913C or WT protein (n=3-5), determined from immunoblots probed with anti-RELN G10 antibodies.

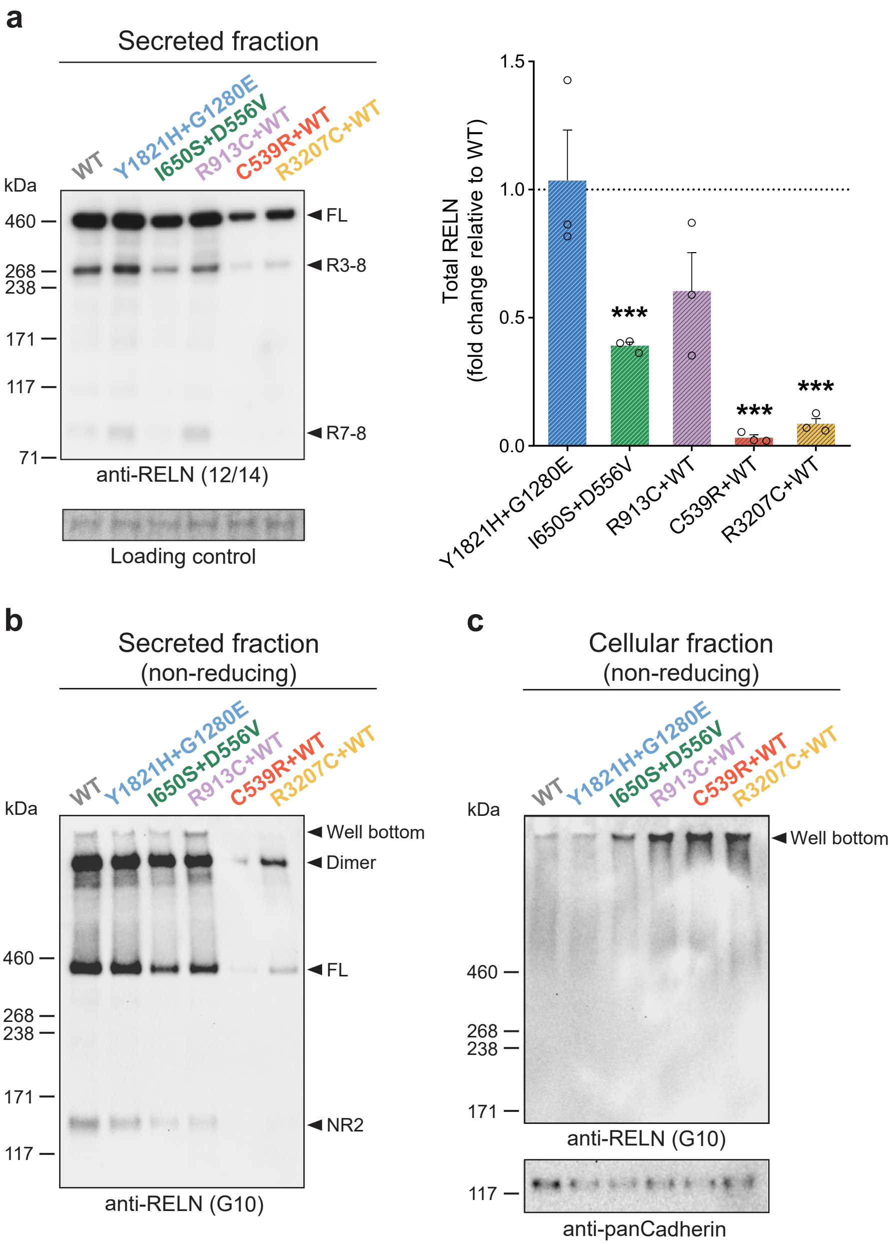


**Supplementary Fig. 2 RELN secretion from co-transfection *in vitro* assay determined with antibodies 12/14 and protein oligomerization.** **a** Representative immunoblotting of the secreted fraction of co-transfected HEK293T cells, probed with anti-RELN 12/14 antibodies. Right panel shows the densitometric analysis of total RELN levels normalized to total protein (Ponceau S) (n=3). Cells transfected with either I650S+D556V, C539R+WT and R3207C+WT presented decreased RELN levels in the media (***$p$=0.0005, ***$p$=0.0001 and ***$p$=0.0005) when compared to WT. **b** Representative immunoblotting of the secreted fraction of co-transfected HEK293T cells in non-reducing conditions (no DTT), probed with anti-RELN G10 antibodies, showing higher bands around 900 kDa and some protein remaining in the well. Strong reduction of any RELN polypeptides and oligomers when co-transfected with the de novo C539R and R3207C mutants supporting their dominant negative effect over the WT protein. **c** RELN in the cellular fraction under non-reducing conditions (no DTT), even when denatured by boiling, failed entering into the gel and remained at the bottom of the well, suggesting that native RELN exists at least as a homodimers intracellularly. An anti-pan Cadherin was used as control. Protein standard sizes (kDa) are indicated on the left side of the blots.

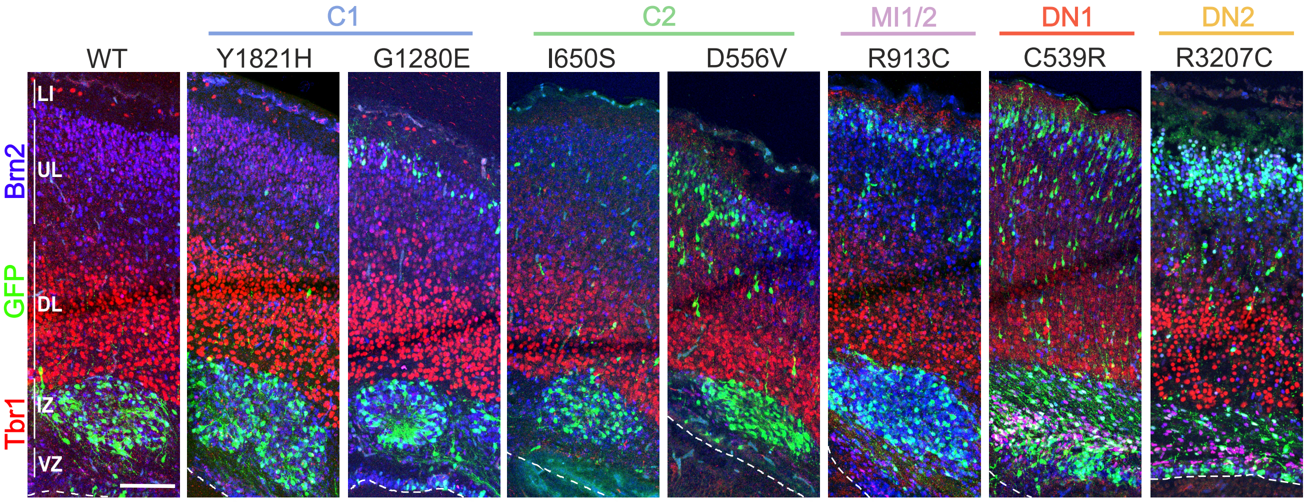


**Supplementary Fig. 3 Aggregates are located in the intermediate zone.** Mosaic maximum projection confocal images showing that in case of aggregate formation, these structures are always localized in the intermediate zone (IZ), immediately under Tbr1$^{+}$ neurons (red) in deep layers although expressing the Brn2 marker of upper layers (blue). For the two mutations C539R and R3207C, which fail to generate aggregates, the electroporated cells (green) that fail to migrate are arrested in the ventricular zone (VZ). Dashed line represents apical limit of VZ. Scale bar: 100 µm.

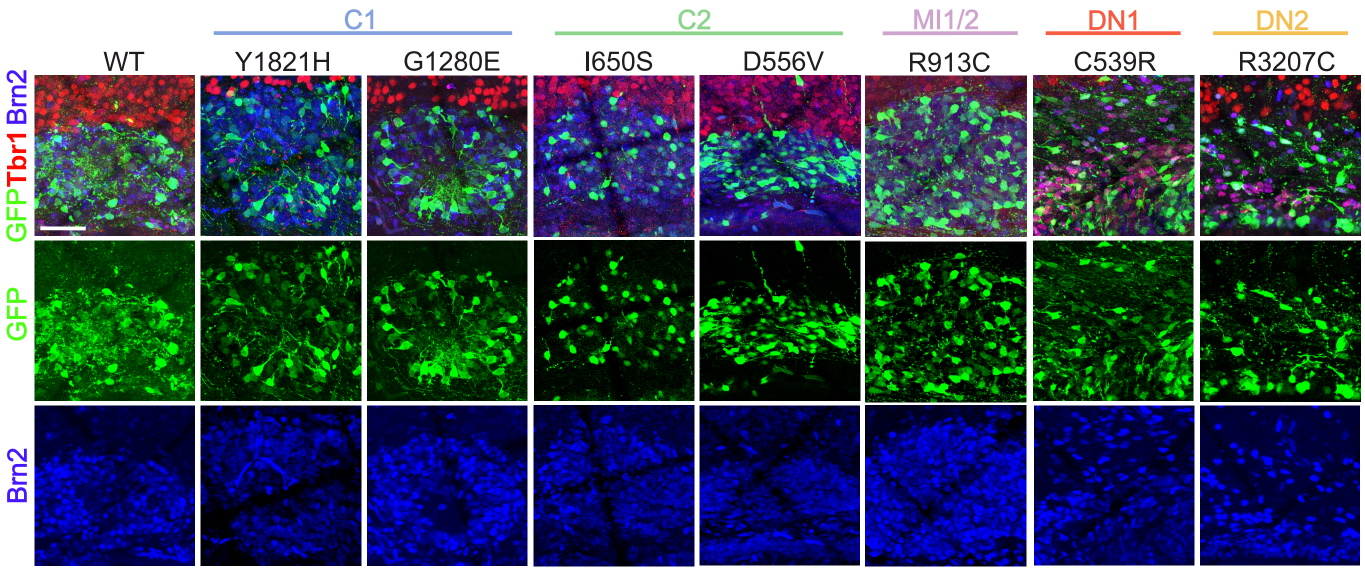


**Supplementary Fig. 4 Fate of neurons forming the rosettes is upper layers.** Mosaic maximum projection confocal images of aggregates. According with the normal birthdating, cells forming the aggregates have an upper layer neuron fate, expressing Brn2 (blue). For mutations C539R and R3207C cells arrested in the VZ are also Brn2$^{+}$. Discontinuities are due to mosaic stitching algorithm. Scale bar: 50 µm.

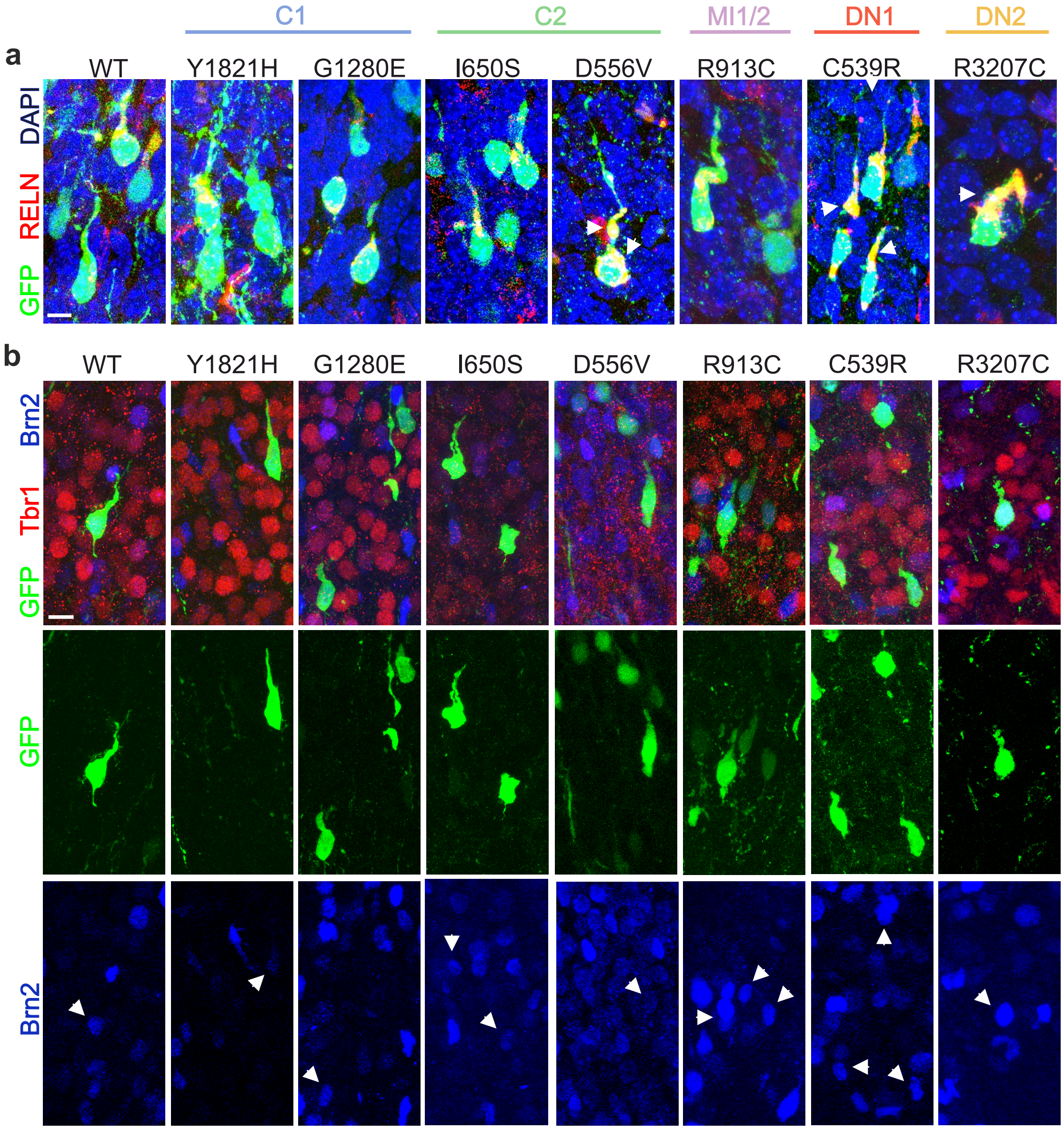


**Supplementary Fig. 5 Representation of glutamatergic neurons delayed in the cortical plate. a** Maximum projection confocal images of $GFP^+$ electroporated neurons delayed in the cortical plate, unable to reach the upper layers at P1. Their morphology is that of still migration neurons and ectopic RELN is accumulated in their apical process. As indicated by the white arrows, cells electroporated with mutations D556V, C539R and R3207C seem to accumulate higher levels of RELN in their cytoplasm. Scale bar: 5 µm. **b** Pyramidal neurons delayed in the cortical plate still keep their fate of upper layer neurons expressing Brn2 (blue, white arrows). Scale bar: 10 µm.

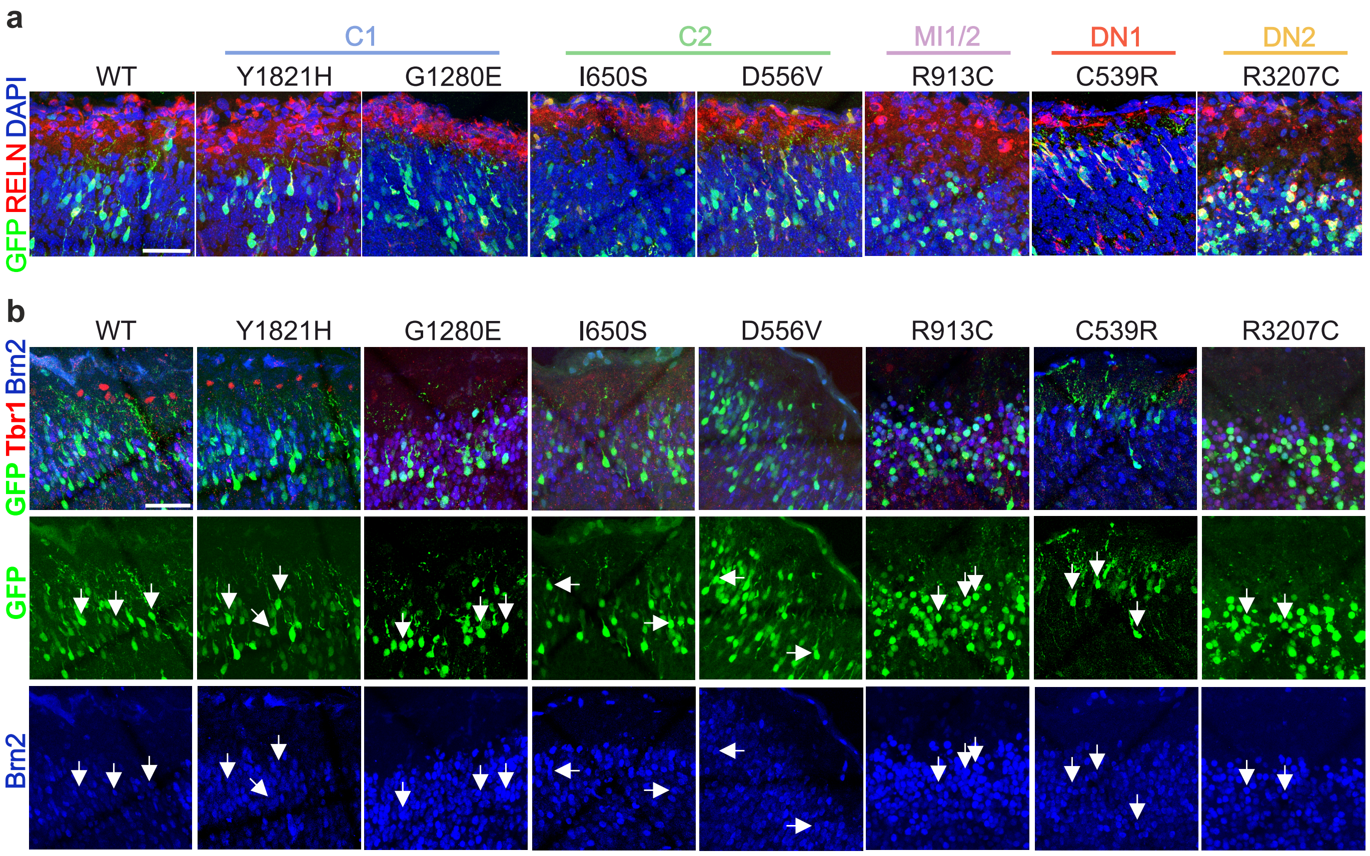


**Supplementary Fig. 6 Electroporated neurons able to reach the upper cortical plate differentiate correctly into upper layers neurons. a** Mosaic maximum projection confocal images of the electroporated cells able to migrate and correctly differentiating into pyramidal neurons directing their dendrites in L1. Scale bar: 50 µm. **b** Mosaic maximum projection confocal images of electroporated neurons reaching the upper layers differentiate correctly Brn2$^{+}$ upper layer neurons (white arrows) as expected by the stage of electroporation. Scale bar: 50 µm.
